# Metabolic modulation of intratumoral cholesterol with gut microbiota for the treatment of colorectal cancer

**DOI:** 10.1101/2025.05.02.651972

**Authors:** Jiayan Wu, Yingyao Lai, Jinming Li, Lixiang Zhai, Pallavi Asthana, Lexin Chen, Minting Chen, Baisen Chen, Susma Gurung, Jin Lyu, Yijing Zhang, Lu Lin, Shuojie Huang, Weihong Kuang, Hui Shi, Yungyung Luo, Xiangxin Kong, Sunny H Wong, Kui Ming Chan, Shuofeng Yuan, Hiu Yee Kwan, Xia Yuan, Weizhen Huang, Yongdui Ruan, Hongmei Lu, Chi Chun Wong, Aiping Lyu, Zhaoxiang Bian, Yixing Ren, Xianjing Wu, Yanlei Ma, Hoi Leong Xavier Wong

**Affiliations:** School of Chinese Medicine, Hong Kong Baptist University, Hong Kong SAR, China; Guangdong Provincial Key Laboratory of Natural Drugs Research and Development, Guangdong Medical University, Dongguan 523808, PR China; Dongguan Key Laboratory of TCM for Prevention and Treatment of Refractory Digestive Diseases, School of Pharmacy, Guangdong Medical University, Dongguan 523808, PR China; Dongguan Key Laboratory of Fundamental Research and Clinical Application of Toxic Chinese Medicine, The First Dongguan Affiliated Hospital, Guangdong Medical University, Dongguan 523710, PR China; Department of Gastroenterology, The First Affiliated Hospital, Guangzhou University of Chinese Medicine, Guangzhou 510405, PR China; Department of Colorectal Surgery, Fudan University Shanghai Cancer Center, Shanghai, China; Department of Oncology, Shanghai Medical College, Fudan University, Shanghai, China; Centre for Chinese Herbal Medicine Drug Development Limited, Hong Kong Baptist University, Hong Kong SAR, China; Center for Obesity and Metabolic Diseases, Gastrointestinal Surgery, Affiliated Hospital of North Sichuan Medical College, Sichuan Province, China; Lee Kong Chian School of Medicine, Nanyang Technological University, Singapore, Singapore; Department of Biomedical Sciences, City University of Hong Kong, Hong Kong SAR, China; State Key Laboratory of Emerging Infectious Diseases, Carol Yu Centre for Infection, Department of Microbiology, School of Clinical Medicine, Li Ka Shing Faculty of Medicine, The University of Hong Kong, Hong Kong SAR, China; Cancer Center of The First Huizhou Affiliated Hospital of Guangdong Medical University, Huizhou 516000, PR China; Department of Acupuncture, The First Dongguan Affiliated Hospital, Guangdong Medical University, Dongguan 523121, PR China; Department of Medicine and Therapeutics, The Chinese University of Hong Kong, Hong Kong SAR, China

**Author notes:** correspondence should be addressed to Wong HLX; Wu X; Ma Y; Ren Y. These authors contributed equally (Jiayan Wu, Yingyao Lai, Jinming Li, Lixiang Zhai, Pallavi Asthana).

## Abstract

Excess cholesterol is positively correlated with colorectal cancer (CRC). Current therapeutic strategies for modulating cholesterol levels in CRC are limited and often come with complications. Here, we demonstrated that microbiome shunting of intestinal cholesterol to anticancer metabolites is a safe and applicable therapeutic approach for CRC. By screening major microbial metabolic products of cholesterol, we found that 4-cholesten-3-one (4-C-3) was selectively depleted in fecal samples and tumor tissues of patients with CRC. 4-C-3 exhibits strong antitumor effects on human CRC cell lines, patient-derived organoids, and patient-derived xenograft (PDX) models. Mechanistically, 4-C-3 suppresses CRC tumorigenesis by dually targeting the epidermal growth factor receptor (EGFR) and Kirsten rat sarcoma viral oncogene homologue (KRAS). 4-C-3 directly binds to EGFR, blocking its signal transduction by inhibiting the binding of EGFR ligands. Additionally, 4-C-3 inhibits oncogenic KRAS variants, such as KRAS^G12D^, by suppressing nucleotide exchange activity and effector engagement. By targeting both EGFR and KRAS, 4-C-3 reduces primary resistance to anti-EGFR therapies caused by KRAS mutations in CRC. As a proof-of-concept study, we showed that delivery of 4-C-3 by *Oscillibacter ruminantium*, a 4-C-3- producing commensal bacterium that is reduced in CRC patients, or a nonpathogenic Escherichia coli strain engineered to specifically convert intestinal cholesterol into 4-C-3, removed intratumoral cholesterol and led to rapid tumor regression in multiple models of CRC. These results suggest that microbial therapies for restoring intestinal cholesterol homeostasis represent a new therapeutic avenue for CRC.

## Introduction

Colorectal cancer (CRC) ranks as the third most common cancer globally, accounting for approximately 1.85 million new cases and 850,000 deaths annually(Siegel et al., 2024). This prevalence emphasizes that CRC represents a significant challenge for healthcare systems worldwide. The treatment of CRC has evolved significantly over the years, but there are still numerous limitations and challenges that affect patient outcomes. CRCs can develop resistance to chemotherapy and other drug treatments, making them less effective and limiting options for patients who have recurrent or metastatic disease. While targeted therapies have shown promise, their efficacy is often limited to patients with specific genetic mutations or biomarkers. A significant proportion of CRC patients may not benefit from these therapies. In addition, a subset of CRC patients, particularly those with high microsatellite instability (MSI-H), may respond well to immunotherapy(Andre et al., 2020; Ganesh et al., 2019; Le et al., 2017). However, the majority of CRC cases are microsatellite stable (MSS) and do not respond as well to these treatments(Andre *et al*., 2020; Chalabi et al., 2020; Ganesh *et al*., 2019; Le *et al*., 2017). In addition, chemotherapy, radiation therapy, and surgery can have significant side effects that can impact the quality of life of patients with CRC. These side effects may also limit the extent of treatment. Therefore, there is an unmet medical need for new therapeutic approaches for CRC.

It has become increasingly clear that microbes residing in the gastrointestinal tract, also known as the gut microbiota, can play a significant role in the development and progression of CRC(Wong and Yu, 2023). Certain pathogenic strains of bacteria, such as *Fusobacterium nucleatum* and *Bacteroides fragilis*, are associated with an increased risk of CRC(Battaglia et al., 2024; Boleij et al., 2015; Bullman et al., 2017; Nejman et al., 2020). The maintenance of a balanced microbial community is therefore crucial for preventing the overgrowth of protumorigenic bacterial species, reducing the risk of inflammation-related carcinogenesis. Probiotics, prebiotics, diet modifications, and fecal microbiota transplants are being studied as ways to modulate the gut microbiota to prevent or treat CRC. Clinical trials and further research are needed to fully understand the mechanisms by which the gut microbiota influences CRC and to translate this knowledge into effective interventions.

Cholesterol is an essential component of cell membranes and a precursor for the synthesis of steroid hormones and bile acids. Despite its importance in physiological functions, altered cholesterol metabolism has been associated with the development of cancers, especially CRC(Kuzu et al., 2016; Pan et al., 2021; Wang et al., 2018). In particular, high serum cholesterol levels are linked to an increased risk of CRC(Fang et al., 2021; Tian et al., 2015). Cholesterol accumulation is a general feature of cancer tissues, including CRC tissues(Cheng et al., 2018). Therefore, cholesterol-lowering strategies have therapeutic potential for CRC prevention and treatment. Some epidemiological studies have suggested that the long-term use of statins, which are commonly used to lower cholesterol, may be associated with a reduced risk of CRC (Cardwell et al., 2014; Nielsen et al., 2012; Voorneveld et al., 2017), but the evidence is not conclusive, and clinical trials have yielded mixed results (Emilsson et al., 2018; Liang et al., 2021). These observations suggest that controlling cholesterol levels alone may not be sufficient to control CRC development in the clinical setting. The long-term use of statins is also associated with diverse complications(Khatiwada and Hong, 2024). Given the limited treatment options for restoring cholesterol homeostasis in cancer, there is an ongoing need for new and safe therapeutic approaches for modulating cholesterol levels in the treatment of CRC. Recent studies suggest that certain species of human gut bacteria, such as *Oscillibacter* spp. and *Eubacterium coprostanoligenes,* can break down cholesterol in the gut, contributing to the reduced risk of cardiovascular diseases in people(Kenny et al., 2020; Li et al., 2024a; Ren et al., 1996). Some bacterial species can also affect the efficiency of cholesterol absorption from the diet(Chen et al., 2018; Le and Yang, 2019). While the gut microbiota can affect cholesterol levels and potentially influence the risk of conditions such as CRC, the relationship is complex and multifactorial. Further research is needed to elucidate the intricate connections among the gut microbiota, cholesterol metabolism, and cancers, especially CRC.

Reducing cholesterol levels via gut microbiota-based interventions is a potential approach for restoring cholesterol homeostasis in CRC. Gut microbiota can promote the conversion of intestinal cholesterol into coprostanol for excretion(Kenny *et al*., 2020; Ren *et al*., 1996). The idea that gut bacteria can control cholesterol levels has been proposed over a century. However, the exact role of this process is still unknown. Despite the longstanding hypothesis, how these interactions might be leveraged for therapeutic purposes in conditions such as hypercholesterolemia and related cancers remains unexplored. In this study, we revealed that engineered microbial therapy aimed at restoring intestinal cholesterol homeostasis is a safe and appliable therapeutic approach for CRC. To explore microbial cholesterol metabolism in CRC, we screened various microbial cholesterol products in the gut and identified that 4-cholesten-3-one (4-C-3) has strong anti-tumor effects in CRC cell lines, patient-derived organoids, and patient-derived xenograft models. We further demonstrated that 4-C-3 is well-tolerated at high doses *in vivo*. Mechanistically, we found that 4-C-3 suppresses CRC tumor growth by targeting both the epidermal growth factor receptor (EGFR) and the Kirsten rat sarcoma viral oncogene homologue (KRAS). This dual targeting allows 4-C-3 to overcome resistance to anti-EGFR therapies caused by KRAS mutations in CRC. Additionally, we engineered a non-pathogenic strain of *Escherichia coli* to convert intestinal cholesterol into 4-C-3. Delivery of 4-C-3 by these bacteria led to rapid tumor regression and reduced intra-tumoral cholesterol in both a chemically induced colitis-associated cancer model and a patient-derived xenograft model, without any signs of toxicity. Clinically, microbial profiling revealed that 4-C-3 and its natural producer, *Oscillibacter ruminantium*, which has anti-tumor properties, are depleted in CRC patients. This proof-of-concept study revealed that engineered microbial therapies are feasible for restoring intestinal cholesterol homeostasis in the treatment of CRC.

## Results

### The level of 4-C-3 reduced in human colon cancer

To address the potential contributions of microbial metabolites of intestinal cholesterol to the development of CRC, we first collected fecal samples from germ-free mice and then detected the contents of microbial products associated with cholesterol metabolism, including 4-cholesten-3-one (4-C-3), 5-cholesten-3-one (5- C-3), coprostanone, and coprostanol(Kenny *et al*., 2020; Ren *et al*., 1996). Consistent with previous studies(Le Roy et al., 2019), fecal cholesterol was significantly enriched in germ-free mice compared with conventional mice **(Fig. S1a)**. However, the above cholesterol metabolites were largely depleted in germ-free mice **(Fig. S1b-e)**, confirming the contribution of the gut microbiota to cholesterol metabolism in the gut. To evaluate the clinical relevance of our findings obtained from mouse studies, the levels of cholesterol and its microbial metabolites were investigated in patients with CRC. In alignment with previous studies(Wu et al., 2022), the increased cholesterol level was observed in CRC tumor biopsies compared with that in adjacent normal tissues (**Fig. 1a**). Among the major microbial metabolic products of cholesterol investigated in this study (**Fig. 1b-e**), we found that 4-C-3 was specifically depleted in tumor biopsies (**Fig. 1c**). We next examined fecal cholesterol and 4-C-3 levels in a CRC patient cohort with 532 samples reported in our previous studies (Kong et al., 2023; Li et al., 2024b; Yang et al., 2021). Consistently, increased fecal cholesterol levels (**Fig. 1f**) and reduced fecal 4-C-3 levels (**Fig. 1g**) were observed in CRC patients compared with healthy controls.

**Figure 1.**
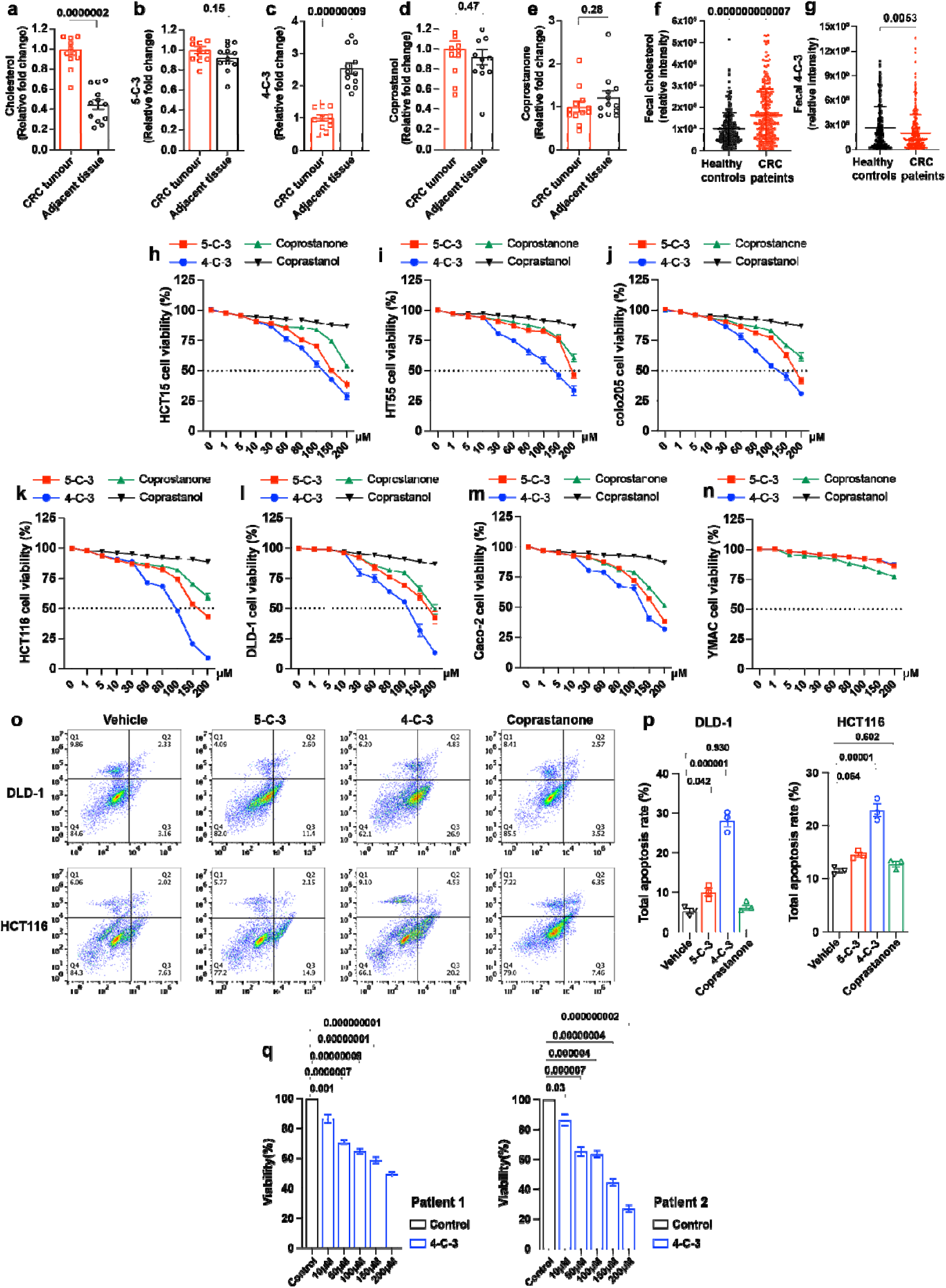
Reduced 4-cholesten-3-one (4-C-3) in human colon cancer and anti-tumor activity of 4-C-3. **(a-e)** Levels of cholesterol and its microbial metabolites 5-cholesten-3-one (5-C-3), 4-cholesten-3-one (4-C- 3), coprostanone, and coprostanol in CRC tumor biopsies and in adjacent normal tissues. (mean ± s.e.m. n=12; Unpaired t test). **(f-g)** Fecal cholesterol and 4-C-3 levels in a CRC patient cohort with 532 samples. (mean ± s.e.m ; Unpaired t test). **(h-n)** MTT assay in a panel of human colorectal cancer cell lines (HCT15, HT55, colo205, HCT116, DLD-1, Caco2 and YMAC) treated with indicated microbial metabolites of cholesterol for 48 h. (mean ± s.e.m., n=3). **(o-p)** Flow cytometry analyses of cell apoptosis in DLD1 and HCT116 cells treated with indicated microbial metabolites of cholesterol for 48 h. The quantification of apoptotic rates is shown in **(p)**. (mean ± s.e.m. n=3; one-way ANOVA). **(q)** Cell viability of colorectal cancer organoids derived from two patients with CRC upon the treatment with 4-C-3 at indicated dosages. (mean ± s.e.m., n=3; one-way ANOVA).

### 4-C-3 protects against colorectal tumorigenesis *in vitro* and *in vivo*

We next investigated the effects of these metabolites in a panel of CRC cell lines, including HCT15, HT55, Colo205, HCT116, DLD-1 and Caco-2 (**Fig 1h-m**). We found that all of these metabolites, except coprostanol, exerted different degrees of inhibitory effects on the growth of CRC cell lines in a dose-dependent manner, whereas no change in growth was observed in the noncancerous colon cell line YMAC (**Fig 1n**), suggesting that the growth-suppressive effects of these metabolites preferentially affect cancer cells. Among the metabolites investigated, 4-C-3, at dosages within the physiological range found in fecal samples of normal human subjects, exhibited the most potent anti-proliferative effect in CRC cell lines (**Fig. 1h-m; S1f).** Consistently, flow cytometry analyses revealed that among the microbial products, 4-C-3 led to the greatest percentage of apoptotic cells in CRC cell lines (**Fig. 1o-p**). The potent anti-CRC effect of 4-C-3 was further confirmed in cancer organoids derived from two CRC patients (**Fig. 1q**). Given the robust anti-CRC effect of 4-C-3 *in vitro* and *ex vivo,* we focused mainly on 4-C-3 in the following *in vivo* investigations.

To confirm the clinical relevance of our findings obtained from the *in vitro* assays, we implanted DLD-1 (**Fig. 2a**) and MC38 (**Fig. S2a)** colon cancer cells subcutaneously into nude mice to generate xenograft tumors. The mice were then gavaged daily with 4-C-3 for 16 days. Treatment with 4-C-3 significantly suppressed DLD-1 (**Fig. 2b-d**) and MC38 (**Fig. S2b-d)** tumor growth in a dose-dependent manner. We next verified that treatment with 4-C-3 suppressed the orthotopic tumor growth of the luciferase-labelled CRC cell line HCT116 (**Fig. 2e**). 4-C-3 significantly reduced tumor size (**Fig. 2f-h**), and intratumoral 4-C-3 levels were markedly increased (**Fig. S3a).** Consistently, a marked decrease in Ki67 levels **(Fig. S3b-c)** and an increase in TUNEL-positive cells **(Fig. S3d-e)** were observed in 4-C-3-treated tumors compared with vehicle-treated controls. To further validate our findings in a more clinically relevant CRC model, we established a patient-derived xenograft (PDX) model by implanting CRC patient-derived cells subcutaneously into nude mice to generate xenograft tumors. The mice were then gavaged daily with 4-C-3 for 15 days (**Fig. 2i**). Treatment with fluorouracil (5-FU) at the optimal therapeutic dosage served as a positive control. Consistently, treatment with 4-C-3 induced a robust protective effect against CRC in mice, which was comparable to that of 5-FU (**Fig. 2j-l**). The 4-C-3 level was consistently significantly increased in the tumors (**Fig. S4)**. Notably, 4-C-3 treatment did not induce any adverse side effect as examined by body weight changes in CRC models (**Fig. S5a-c)**. Furthermore 4-C-3 did not alter normal tissue homeostasis, as assessed by body weight (**Fig. S5d)** and regular blood tests (**Fig. S5e-f)**. These *in vitro* and *in vivo* data, with the support of multiple tumor models, demonstrate a safe and potent effect of 4-C-3 in modulating CRC growth.

**Figure 2.**
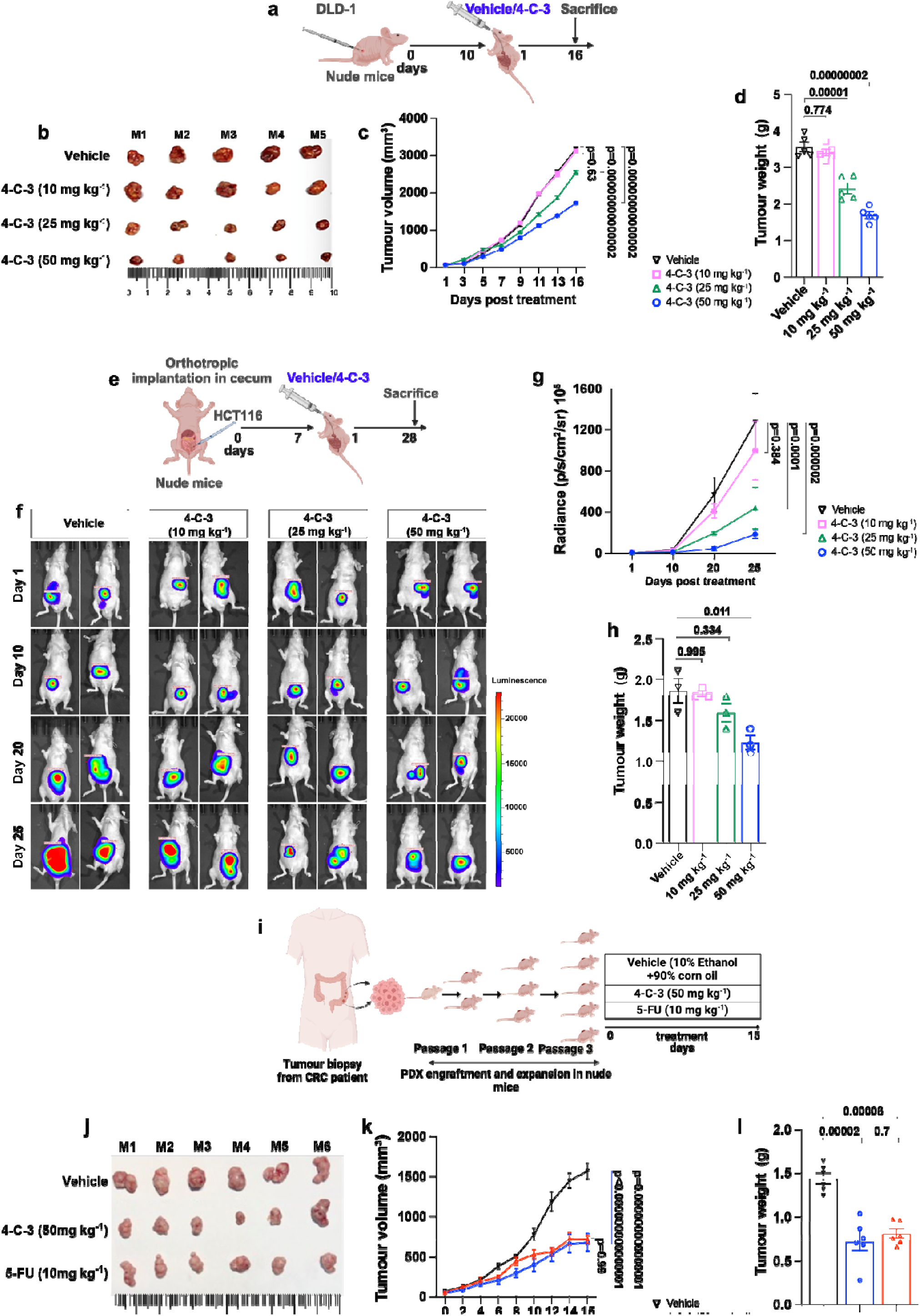
4-cholesten-3-one (4-C-3) suppresses colon tumorigenesis *in vivo*. **[a-d] (a)** Schematic representation demonstrating the experimental design for the establishment of subcutaneous CRC model with DLD-1 cells along with the treatment with 4-C-3 at indicated dosages. **(b)** Images of DLD-1 xenograft tumors upon the treatment with 4-C-3. Tumor volume **(c)** and tumor volume **(d)** of DLD-1 xenograft tumors of mice in the indicated groups. [mean ± s.e.m., n=5 for each treatment group; two-way ANOVA for **(c)** & one-way ANOVA for **(d)**. **[e-h] (e)** Schematic representation showing the experimental design for establishing orthotropic CRC model with luciferase-labelled HCT116 cells and the treatment with 4-C-3 at indicated dosages. **(f)** Images for luciferase signal of HCT116 xenograft tumors after the treatment with 4-C-3. **(g)** Quantification of luciferase signal emitted by HCT116 xenograft tumors for the measurement of tumor growth. **(h)** Tumor weight. [mean ± s.e.m., n=3 for each treatment group; two- way ANOVA for **(g)** & one-way ANOVA for **(h)**. **[i-l] (i)** Schematic diagram demonstrating the experimental design for the establishment of subcutaneous CRC model with patient-derived cancer cells (PDX) and indicated treatment groups. **(j)** Images of PDX xenograft tumors upon the indicated treatments. **(k-l)** Tumor volume **(k)** and tumor weight **(l)** of PDX xenografts in the indicated groups. [mean ± s.e.m., n=6 for each treatment group; two-way ANOVA for **(k)** & one-way ANOVA for **(l)**]

### 4-C-3 inhibits the growth of CRC cells by antagonizing the EGFR pathway

In our efforts to provide molecular and mechanistic insights into the anti-CRC effects of 4-C-3, we employed transcriptomics to determine the gene modules regulated by 4-C-3 in CRC cells. RNA sequencing studies in DLD1 cells treated with 4-C-3 revealed dramatic transcriptional changes induced by 4-C-3 (**Fig. 3a**). Volcano plot analysis demonstrated that 129 and 326 genes were up-regulated and down-regulated by 4-C-3, respectively. KEGG pathway enrichment analysis of down-regulated genes identified the EGFR signaling pathway as the top enriched pathway (**Fig. 3b**). Gene set enrichment analysis (GSEA) confirmed suppressed EGFR signaling in 4-C-3-treated cells (**Fig. 3c**). Consistently, qPCR analyses validated that the expression of epiregulin (EREG), a ligand of EGFR that is highly expressed in CRC tumors and regulated by an autocrine loop through EGFR downstream signaling activation(Cheng et al., 2021; Qu et al., 2016), was significantly reduced in DLD-1 cells treated with 4-C-3 (**Fig. 3d**). As a complementary approach, we observed that treatment with 4-C-3 significantly inhibited the phosphorylation of EGFR and its downstream signaling targets, including PI3K, Akt and m-TOR, in the DLD-1 cell line (**Fig. 3e**). A similar observation was obtained for orthotopic DLD1 tumors from 4-C-3-treated mice (**Fig. 3f**). The inhibitory effect of 4-C-3 on the growth of Caco-2 cell line was abrogated by EGFR ablation with siRNA (**Fig. 3g**), confirming that EGFR signaling is an intermediary of the inhibitory effect of 4-C-3 on CRC growth. To elucidate the mechanism of EGFR inhibition by 4-C-3, we investigated the potential interaction between EGFR and 4-C-3 via a cellular thermal shift assay in HCT116 and DLD-1 cell lines. In relative to the control, EGFR exhibited thermal shifts in the presence of 4-C-3 at the denaturation temperate ranging from 51°C to 57°C, revealing the cell-level interaction between EGFR and 4-C-3 (**Fig. 3h**). We then performed microscale thermophoresis (MST) analyses on binding between the recombinant extracellular domain of EGFR (EGFR-ECD) and 4-C-3. EREG, one of the top downregulated targets by 4-C-3 in our transcriptome analyses (**Fig. 3a**), acted as a positive control for the assay. We established that 4-C-3 directly bound to the recombinant extracellular domain of EGFR with a K_D_ value of 5.6 µM while EREG had a K_D_ value of 0.5 µM (**Fig. 3i-j**). Although EREG bound EGFR-ECD with higher affinity, the binding of EREG was completely abrogated when EGFR-ECD was co-incubated with EREG and 4-C-3 (**Fig. 3k**). By performing immunoprecipitation with Caco2 cells, we confirmed that 4-C-3 efficiently inhibited the cell-surface binding of EREG to EGFR (**Fig. 3l**), revealing the ability of 4-C-3 to block the ligand binding to EGFR. To further validate this observation, we performed additional MST experiments with a high affinity ligand EGF and a low affinity ligand AREG. 4-C-3 similarly suppressed the bindings of EGF and AREG to EGFR-ECD **(Fig. S6a-b)**. To determine the residues of EGFR involved in binding with 4-C-3, we docked 4-C-3 into an AlphaFold-modelled EGFR. The binding pocket of EGFR-ECD around 4-C-3 was calculated to be at a distance of 3.1 Å. In the modelled binding pocket, 4-C-3 interacts with Asn528 in the EGFR binding pocket via H-bond formation (**Fig. 3m**). To further understand how 4-C-3 interrupts ligand binding to EGFR, we executed 100ns molecular dynamics (MD) simulations coupled with mechanics and thermodynamic calculations to study the dynamical structural characteristics and interactions between them **(Fig. S6c-f)**. During the MD simulation, EREG was initially placed in the binding site of EGFR at a distance of 4 Å with a binding free energy (vdW=-75.39kcal/mol; Coulomb=-31.02kcal/mol) **(Fig. S6c)**. This binding was stabilized by multiple non-bonded interactions, including hydrogen bonding, salt bridge interactions, π-π stacking interactions and π- cation interactions **(Fig. S6d)**. Upon the binding of 4-C-3, EGFR underwent gradual conformational changes, leading to the destabilization of EREG/EGFR complex **(Fig. S6e)**. By 90ns, EREG was completely dissociated from EGFR as evidenced by the increased binding distance (36 Å), the reduced binding energy (vdW=0 kcal/mol; Coulomb=-8kcal/mol) and the disruption of all non-bonded interactions **(Fig. S6f)**. Taken together, these findings reveal that 4-C-3 is a potent endogenous inhibitor of EGFR and that it inhibits the growth of CRC cells by antagonizing the EGFR pathway.

**Figure 3.**
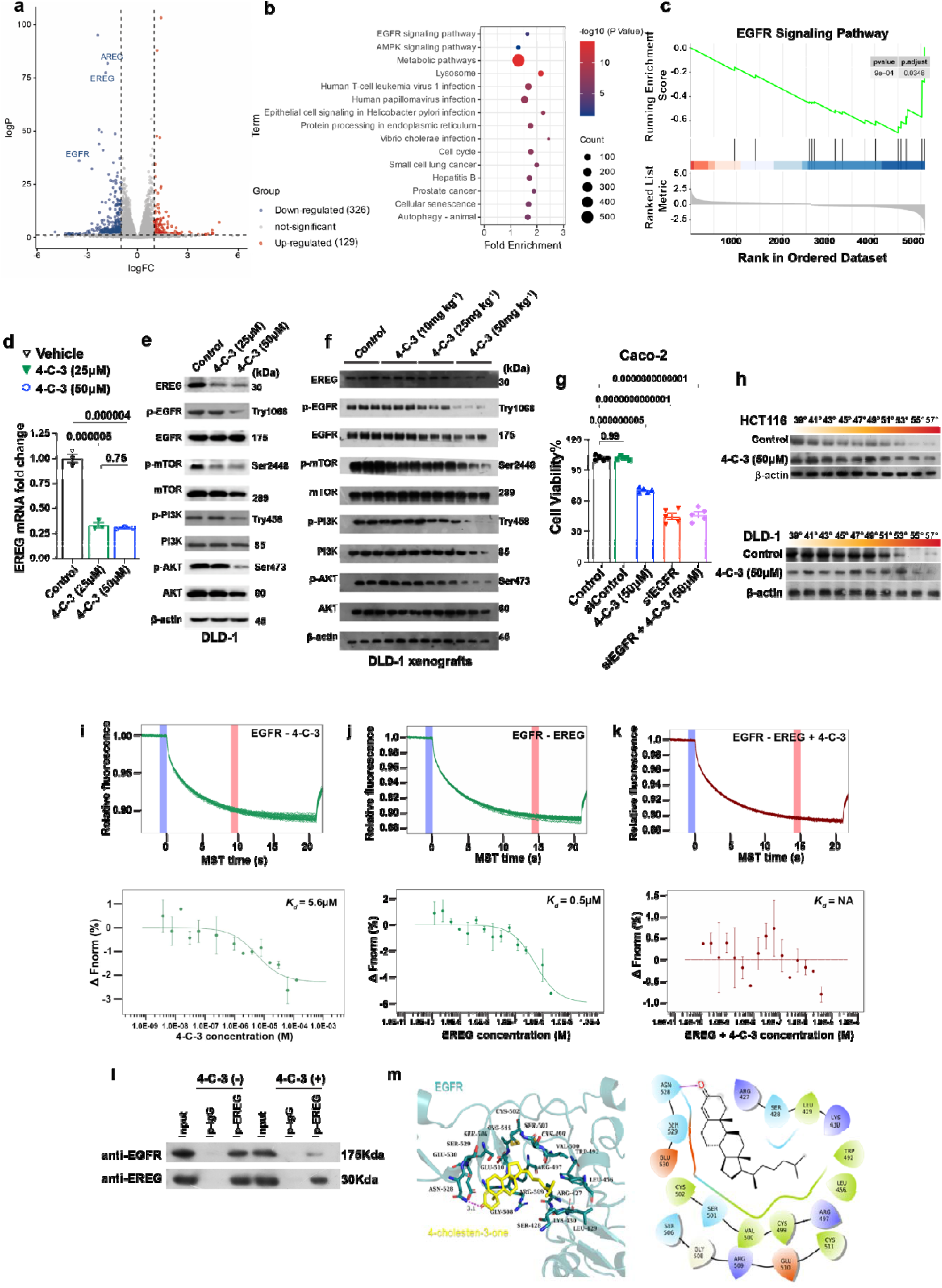
4-cholesten-3-one (4-C-3) suppresses colon tumorigenesis through suppression of EGFR signaling. **(a-c)** RNA sequencing analyses of DLD-1 cells treated with 4-C-3 (50µM) for 48h. Volcano plot analysis **(a)**, KEGG pathway enrichment analysis **(b)** and gene set enrichment analysis **(c)** were performed. **(d)** qPCR analysis of the expression of EREG in DLD-1 cells treated with 4-C-3 at indicated dosages for 48h. (mean ± s.e.m., n=3; one-way ANOVA). **(e-f)** Western blotting analyses of EGFR signaling pathway in *in vitro* DLD-1 cells treated with 4-C-3 at indicated dosages **(e)** (n=3) and DLD-1 subcutaneous xenografts upon the treatment with 4-C-3 at indicated dosages **(f)** (n=3 for each treatment group). **(g)** MTT assay of Caco-2 cells with siRNA-mediated knockdown of EGFR in response to the treatment with 4-C-3 at the indicated dosage (mean ± s.e.m., n=3; one-way ANOVA). **(h)** Representative images for western blotting analyses for cellular thermal shift assay of EGFR in HCT116 and DLD-1 cells upon the treatment with 4-C- 3 (50µM). (n=3). **(i-k)** Microscale thermophoresis (MST) analyses for the binding between the recombinant extracellular domain of EGFR (EGFR-ECD) and 4-C-3 **(i)**; the binding between EGFR-ECD and recombinant EREG **(j)** and the binding between EGFR-ECD and recombinant EREG in the presence of 4-C- 3 **(k)**. (n=3)**. (l)** Co-immunoprecipitation of EGEG and EGFR in lysates of DLD-1 cells treated with 4-C-3, followed by western blotting using antibodies indicated. (n=3). **(m)** Molecular docking analysis of the binding between EGFR and 4-C-3.

### 4-C-3 attenuates resistance to common anti-EGFR therapies in CRC

Anti-EGFR therapies, such as cetuximab and panitumumab, are effective therapeutic agents for metastatic CRC(Martinelli et al., 2020). However, resistance to targeted therapies limits their clinical use and efficiency(Martinelli *et al*., 2020). To investigate whether 4-C-3 delivery can attenuate resistance to common anti-EGFR therapies in CRC, the inhibitory effect of 4-C-3 on CRC growth was examined in Caco- 2 cells expressing wild-type EGFR or mutant EGFRs with common drug resistance mutations (EGFR^S492R^ and EGFR^G719S^). Treatment with cetuximab served as a control. Unlike Caco-2 cells with wild-type EGFR, cells expressing mutant EGFRs were all resistant to cetuximab at our tested dosages (**Fig. 4a**). In contrast, 4- C-3 at the same molar dose effectively suppressed the growth of these EGFR-mutant cells (**Fig. 4a**), suggesting that 4-C-3 attenuates resistance to common EGFR resistance mutations. Furthermore, molecular docking analysis showed that 4-C-3 interacts with an epitope on EGFR that is distinct from the cetuximab epitope (**Fig. S7a-b)**, revealing the potential mechanism by which 4-C-3 overcomes cetuximab resistance in CRC. In colorectal cancer, the presence of KRAS mutations is a significant predictor of resistance to epidermal growth factor receptor (EGFR) inhibitors.(Cerami et al., 2012; Gao et al., 2013; Zehir et al., 2017). These mutations, often found in codons 12 and 13 of the *KRAS* gene, lead to constitutive activation of the RAS/RAF/MEK/ERK signaling pathway, rendering EGFR-targeted therapies ineffective(Liu et al., 2019; Modest et al., 2016). In contrast, agents targeting KRAS mutations have limited efficacy against KRAS-mutant CRC in part because of EGFR-mediated reactivation of MAPK signaling, prompting clinical trials with combined therapies targeting KRAS mutations and EGFR in CRC. However, secondary resistance mechanisms driven by the compensatory activation of other tyrosine kinases or the upregulation of mTOR signaling limit the response to the combination strategy(Ryan et al., 2022; Whitehead et al., 2024; Yaeger et al., 2023). Given the EGFR-PI3K-mTOR-inhibitory profile of 4-C-3, we postulated that 4-C-3 may be efficacious against KRAS-mutant CRC. We indeed found that 4-C-3 also had potent inhibitory effects on the proliferation of KRAS-mutant CRC cell lines, including LS513(KRAS^G12D^) and HCT116 (KRAS^G13D^), both of which were resistant to cetuximab (**Fig. 4b**). While cetuximab was able to decrease EGFR activation in LS513 and HCT116 cells, the cells maintained a high level of PI3K/mTOR signaling activation that was not suppressed by drug treatment. In contrast, 4-C-3 effectively inactivated the phosphorylation of EGFR and its downstream signaling targets (m-TOR, PI3K, AKT and Erk1/2) in these cell lines (**Fig. 4c**). To validate the anti-tumor efficacy of 4-C-3 on KRAS mutant CRC *in vivo,* we treated nude mice bearing subcutaneous LS513 or HCT116 xenografts with either cetuximab or 4-C-3. As expected, 4-C-3 conferred potent inhibitory effects on the growth of LS513 and HCT116 xenografts, whereas cetuximab had limited effects (**Fig. 4d-i**). In addition, we further tested the anti-tumor effect of 4-C-3 on CRC organoids derived from *Apc^min/+^Kras^G12D/+^* mutant mice. Treatment with 4-C-3 suppressed the growth of mutant Kras-CRC organoids, with potency comparable to that of the KRAS^G12D^ inhibitor MRTX1133 (**Fig. 4j**). Consistently, western blot analysis showed that 4-C-3 inhibited EGFR-MAPK signaling activation in these organoids (**Fig. 4k**). These findings demonstrate that 4-C-3 holds promise for overcoming mutant KRAS-driven resistance to anti-EGFR therapies in the treatment of CRC.

**Figure 4.**
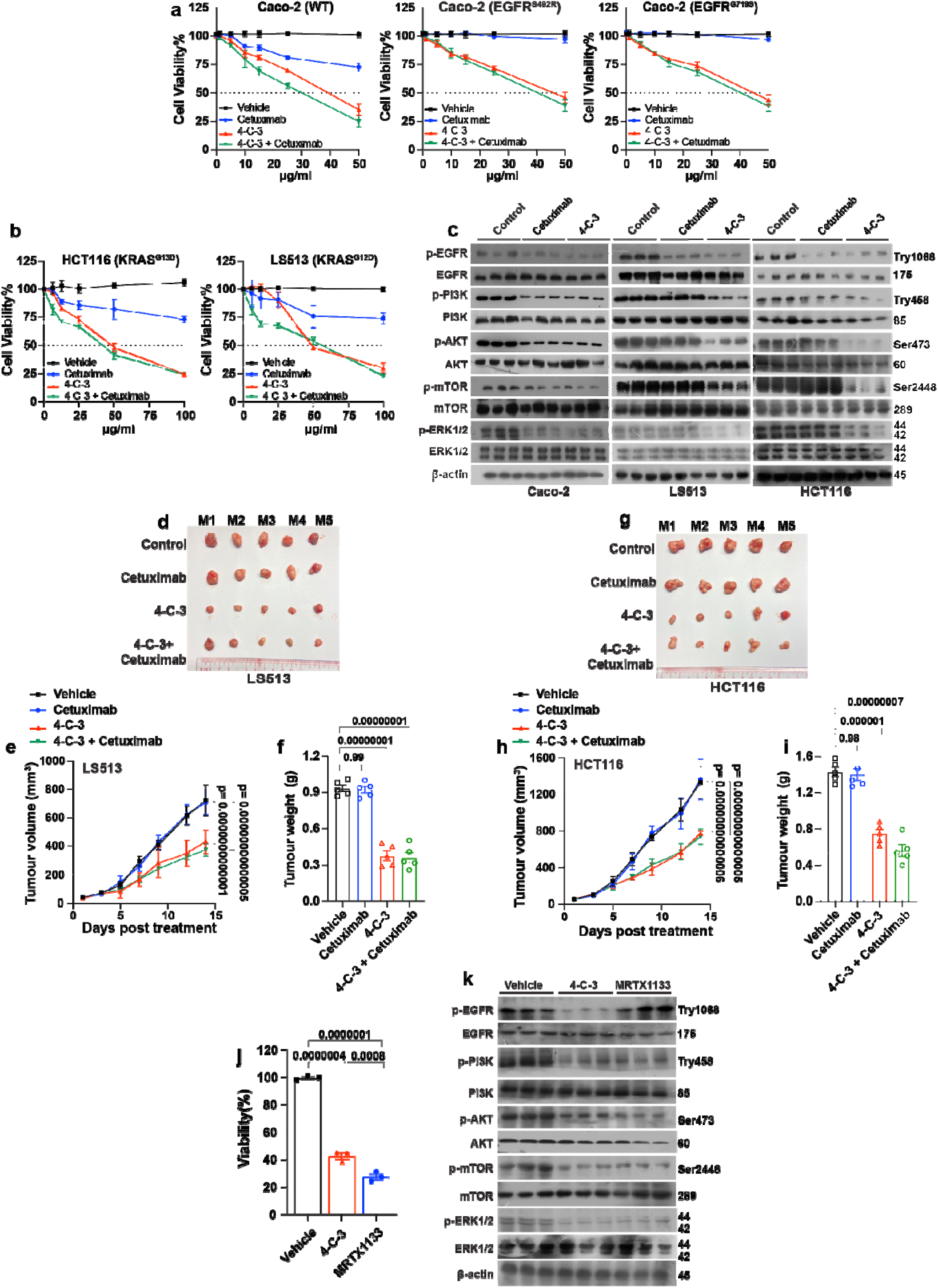
4-cholesten-3-one (4-C-3) overcomes primary resistance to anti-EGFR therapeutics in C C. **(a)** MTT assay of Caco-2 cells expressing wild-type EGFR and indicated EGFR mutants treated with either 4-C-3 and cetuximab. (mean ± s.e.m., n=3). **(b)** MTT assay of HCT116 and LS513 cells in response to the treatment with 4-C-3 or cetuximab. (mean ± s.e.m., n=3). **(c)** Western blotting analyses of EGFR and its downstream signaling pathways in Caco-2, LS513, and HCT116 cells upon the treatment with 4-C-3 or cetuximab (n=3). **[d-i]** Nude mice bearing subcutaneous LS513 or HCT116 xenografts were treated with either cetuximab or 4-C-3. **(d, g)** Images of tumors for indicated groups are shown. **(e-f)** Tumor volume **(e)** and tumor weight **(f)** of LS513 xenografts. **(h-i)** Tumor volume **(h)** and tumor weight **(i)** of HCT116 xenografts. [mean ± s.e.m., n=5 for each treatment group; two-way ANOVA for **(e & h)** & one-way ANOVA for **(f & i)**]. **(j)** Cell viability of CRC organoids derived from *Apc^min/+^Kras^G12D/+^* mutant mice upon the treatment with 4-C-3 and KRAS^G12D^ inhibitor MRTX1133. (mean ± s.e.m., n=3; one-way ANOVA). **(k)** Western blotting analyses of EGFR and its downstream signaling pathways in CRC organoids upon the treatment with 4-C-3 or MRTX1133 (n=3).

### 4-C-3 is a non-covalent small molecule inhibitor of KRAS oncogenic variants

Our observations that 4-C-3 is efficacious against multiple KRAS-mutant CRC cells are surprising, given the expectation that KRAS oncogenic mutations are causally implicated in resistance to EGFR blockage in CRC, and the inhibitory effect of 4-C-3 on EGFR activation is theoretically not sufficient to attenuate the growth of KRAS-mutant CRC cells. This prompted us to explore additional mechanisms that may contribute to the ability of 4-C-3 to target KRAS-mutant CRC cells. As the combinatorial targeting of EGFR and KRAS is an efficacious therapeutic strategy to treat patients with KRAS mutation-driven CRC, we postulated that KRAS may be a possible target for 4-C-3 in CRC. In this study, we focused on the KRAS p.G12D for investigation, as the *KRAS^G12D^* mutation is the most common KRAS mutation in CRC, and there is currently no FDA-approved drug for targeting cancers with this mutation. Using MST assays, we found that 4-C-3 exhibited 3-fold selectivity for binding to GDP-bound KRAS^G12D^ compared with GDP-bound KRAS^WT^ (GDP-bound KRAS^G12D^ K_D_= 7.2µM versus GDP-bound KRAS^WT^ K_D_= 21.3µM) (**Fig. 5a**). To examine the potential effects of 4-C-3 on KRAS functions, fluorescent-based biochemical assays were employed to study the rate of GDP/GTP exchange reactions of KRAS. 4-C-3 reduced the rates of both intrinsic and SOS1-mediated nucleotide exchange reactions for KRAS^G12D^, with IC_50_ values of 14.32µM and 12.57µM, respectively (**Fig. 5b-c**). For KRAS^WT^, 4-C-3 had weaker effects, with IC50 values of 30.79µM and 28.4µM, respectively (**Fig. 5b-c**). This indicates that 4-C-3 exerts stronger inhibition on KRAS^G12D^ than on KRAS^WT^. To investigate the effect of 4-C-3 on effector binding, we tested the potency of 4-C-3 to displace the RAS-binding domain of CRAF from purified KRAS variants preloaded with GMPPNP, a non-hydrolysable GTP analog. 4-C-3 diminished the active KRAS^G12D^-CRAF interaction with an IC_50_ value of 41.14µM, but did not affect the interaction between CRAF and KRAS^WT^ at this concentration (**Fig. 5d**), which again suggests that 4-C-3 is more selective for KRAS^G12D^. To further confirm the interaction between 4-C-3 and KRAS at the cellular level, we performed traditional cellular thermal shift assays in CaCO2 (KRAS^WT^) and LS513 (KRAS^G12D^) cell lines. Compared to the control, KRAS exhibited significant thermal shifts in the presence of 4-C-3 in LS513 cells, whereas this thermal shift was less prominent in CaCO2 cells (**Fig. 5e**), revealing the intracellular interaction between KRAS and 4-C-3 and the selectivity of 4-C-3 towards KRAS^G12D^ binding. To decipher the regulatory role of 4-C-3 in KRAS signaling, we performed an active RAS (RAS-GDP) pull-down assay in the cell lysate of LS513 cells treated with 4-C-3. Our results showed that 4-C-3 reduced the level of KRAS-GTP in LS513 cells (**Fig. 5f**). Additionally, 4-C-3 effectively suppressed the phosphorylation of Erk1/2, a key downstream pathway in KRAS signaling, in these cells, whereas cetuximab had limited effects (**Fig. 4c**). To better understand how 4-C-3 interacts with KRAS, we docked 4-C-3 into an AlphaFold-modelled GDP-bound KRAS^G12D^. 4-C-3 was predicted to bind in the switch-II pocket of KRAS though H-bond formation with Asp12, an interaction that could not be observed in KRAS^WT^ (**Fig. 5g**). We next performed long timescale simulations to reveal the interaction dynamics of the 4-C-3-KRAS^G12D^ complex **(Fig. S8a-d)**. The protein backbone root-mean-square fluctuation (RMSF) values showed that the KRAS^G12D^-GDP-4-C-3 system reached equilibrium at 8ns, while the KRAS^G12D^- GDP system in the absence of 4-C-3 did not reach equilibrium within 30ns **(Fig. S8a-b)**. This indicates that 4-C-3 stabilizes the KRAS^G12D^-GDP complex. In contrast, the RMSD values for the KRAS^WT^-GDP-4-C-3 and KRAS^WT^-GDP systems showed that equilibrium was reached at 12 ns with 4-C-3 and at 16 ns without it **(Fig. S8c-d)**. This suggests that 4-C-3 only slightly stabilizes the KRAS^WT^-GDP complex. To validate these simulations, we performed MST assays on the binding of GDP on KRAS^G12D^. Upon the addition of 4-C-3, the binding between GDP and KRAS^G12D^ increased by approximately 6-fold **(Fig. S8e-f)**. These results suggest that the binding of 4-C-3 locks KRAS^G12D^ in an inactive, GDP-bound state. These results collectively suggest that the dual-targeting of EGFR and KRAS with 4-C-3 is an effective therapeutic strategy for overcoming mutant KRAS-driven resistance to anti-EGFR therapies in CRC.

**Figure 5.**
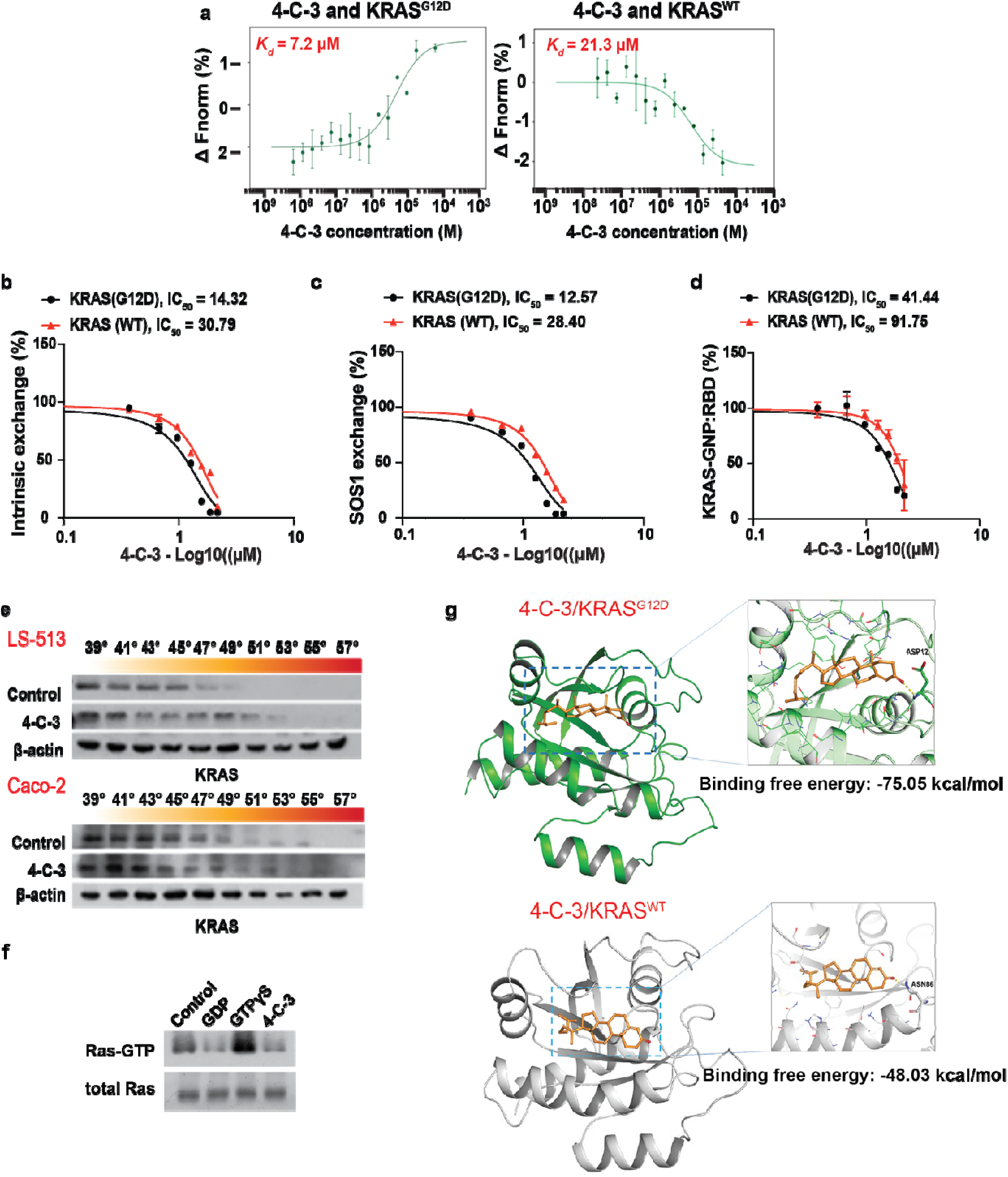
4-cholesten-3-one (4-C-3) is a non-covalent inhibitor for oncogenic KRAS mutants. **(a)** Microscale thermophoresis (MST) analyses for the binding between recombinant KRAS^WT^/KRAS^G12D^ bound with GDP and 4-C-3. (n=3). **(b-c)** The effect of 4-C-3 treatment on the intrinsic **(b)** and SOS1 **(c)**- mediated nucleotide exchange in KRAS^WT^/KRAS^G12D^. (n=3) **(d)** KRAS^WT^/KRAS^G12D^ were pre-loaded with the non-hydrolyzable GTP analogue GMPPNP (GNP), followed by the reaction with a GST-tagged RBD domain of CRAF. The mixtures were then analyzed by GST pull-down and immunoblotting for quantification with densitometry. (n=3). **(e)** Representative images for western blotting analyses for cellular thermal shift assay of KRAS in LS513 and Caco-2 cells upon the treatment with 4-C-3 (50µM). (n=3). **(f)** LS513 cells, a KRAS G12D mutant cell line, were treated with indicated treatments and their extracts were subjected to RBD pull-down and immunoblotting to determine the level of active KRAS (KRAS-GTP). (n=3). **(g)** Molecular docking analysis of the binding between KRAS^WT^/KRAS^G12D^ and 4-C-3.

### The development of functional engineered commensal microbes for the treatment of CRC

In solid tumors, especially CRC, high levels of cholesterol are often observed; this phenomenon has been linked to increased tumor proliferation and progression (Kuzu *et al*., 2016; Pan *et al*., 2021; Wang *et al*., 2018). Given the potent inhibitory effect of the cholesterol metabolite 4-C-3 on CRC growth, we next investigated whether microbial catabolism of cholesterol into 4-C-3 by the gut microbiota can inhibit CRC tumorigenesis. A previous study revealed that the cholesterol oxidase (CHOX) from marine *Streptomyces* sp. can specifically catalyze the oxidation and isomerization of cholesterol to 4-C-3 (Ren *et al*., 1996). To facilitate the conversion of cholesterol to 4-C-3 in the tumor, we transformed a series of nonpathogenic strains of *E. coli* including *E. coli*. (TOP 10), *E. coli*. (JM109) and *E. coli*. (Nissle 1917) with a single plasmid that encodes CHOX under the control of the constitutive promoter P194 (**Fig. S9a)**. In *in vitro* culture with a medium containing cholesterol, we found that only the engineered *E. coli*. (TOP 10) bacteria expressing CHOX could efficiently convert cholesterol to 4-C-3, as assessed by liquid chromatography tandem mass spectrometry (LC MS) analysis (**Fig. S9b-c)**. This 4-C-3-producing strain is hereafter referred to as *E. coli CHOX^+^* bacteria. To study the *in vivo* capacity of engineered *E. coli CHOX^+^* bacteria in producing 4-C-3, we subcutaneously implanted MC38 colon cancer cells into nude mice to generate xenograft tumors. The mice were then gavaged daily with *E. coli CHOX^+^* bacteria or *E. coli CHOX^-^* bacteria. The colonization of *E. coli CHOX^+^*bacteria but not *E. coli CHOX^-^* bacteria significantly reduced MC38 tumor growth after treatment initiation (**Fig. S10a-c)**. The level of 4-C-3 was significantly increased in treated tumors (**Fig. S10d)**, confirming the successful delivery of 4-C-3 to the tumor by our engineered bacteria. Notably, the colonization of *E. coli CHOX^+^* bacteria exhibited an excellent safety profile in this cancer model, as assessed by body weight (**Fig. S11a)**. Furthermore *E. coli CHOX^+^* bacteria showed no adverse effects in healthy mice as shown by body weight and regular blood tests results (**Fig. S11b-d)**.

### *Oscillibacter ruminantium-*mediated cholesterol metabolism protects against CRC tumorigenesis

A recent study revealed that some *Oscillibacter* species, culturable gut bacteria with relatively high levels in the human gut microbiome, can metabolize cholesterol and produce its metabolites, including 4-C-3 (Li *et al*., 2024a). To investigate whether these bacterial species are responsible for cholesterol dysregulation in CRC, we analyzed our previously published datasets obtained from metagenomic analyses in our CRC patient cohort (Kong *et al*., 2023; Li *et al*., 2024b; Yang *et al*., 2021) and identified that *Oscillibacter ruminantium (O. ruminantium)* could convert cholesterol into 4-C-3 in *in vitro* culture (**Fig. S12)**. Unlike *E. coli CHOX^+^* bacteria, which specifically convert cholesterol into 4-C-3, *O. ruminantium* produces both 4-C- 3 and its downstream metabolites. This explains the higher capacity of *E. coli CHOX^+^* bacteria in producing 4-C-3 (**Fig. S12)**. The abundance of *O. ruminantium* was reduced in CRC patients when compared with healthy controls (**Fig. 6a**). To investigate whether the conversion of cholesterol into 4-C-3 by *O. ruminantium* similarly exerts anti-tumor effects on CRC as observed with *E. coli CHOX^+^* bacteria, we daily gavaged mice bearing orthotropic HCT116 xenograft tumors with *O. ruminantium* (DSM113516)*, E. coli CHOX^+^* bacteria or 4-C-3 for 21 days. *Oscillibacter valericigenes (O. valericigenes)*, which could not produce 4-C-3, served as a control. All treatment groups, except for the colonization with *O. valericigenes,* signficantly suppressed tumor growth, with the strongest effects seen in mice colonized with *E. coli CHOX^+^*bacteria (**Fig. 6b-d**), which correlated with the higher capacity of *E. coli CHOX^+^* bacteria to convert cholesterol into 4-C-3 (**Fig. 6e-f**). Consistently, similar results were obtained in a murine model of CRC induced with a combination of the carcinogen azoxymethane (AOM) and colitis-inducing dextran sodium sulfate salt (DSS) over a period of 14 weeks (**Fig. 6g**). All treatment groups, except for *O. valericigenes*, increased colon length, and reduced tumor load and number, with the most robust effect observed in the colonization of *E. coli CHOX^+^* bacteria (**Fig. 6h-j**). Along with the reduction in intratumoral cholesterol (**Fig.6j)**, the level of 4-C-3 increased significantly in the tumors treated with *O. ruminantium or E. coli CHOX^+^* bacteria (**Fig. 6k**). Next, we investigated the ability of *Oscillibacter ruminantium* to mitigate mutant KRAS-induced CRC tumorigenesis. CRC organoids from *Apc^min/+^Kras^G12D/+^* mutant mice were subcutanously implanted in immunocompetent mice. The mice were subjected to a 10-day regimen with either daily intratumoral injection of *O. ruminantium* (DSM113516) or *E. coli CHOX^+^* bacteria, or oral administration of 4-C-3, or intraperitoneal administration of the KRAS^G12D^ inhibitor MRTX1133. Again, all treatment groups provoked anti-tumor effects on Kras^G12D^ CRC tumors, with the efficacy of *E. coli CHOX^+^*bacteria comparable to that of MRTX1133 (**Fig. 6l-p**). Collectively, these data suggest that microbiome-mediated shifting of the balance from protumor cholesterol to antitumor 4-C-3 in the intratumoral environment may be an applicable therapeutic approach for the treatment of CRC.

**Figure 6.**
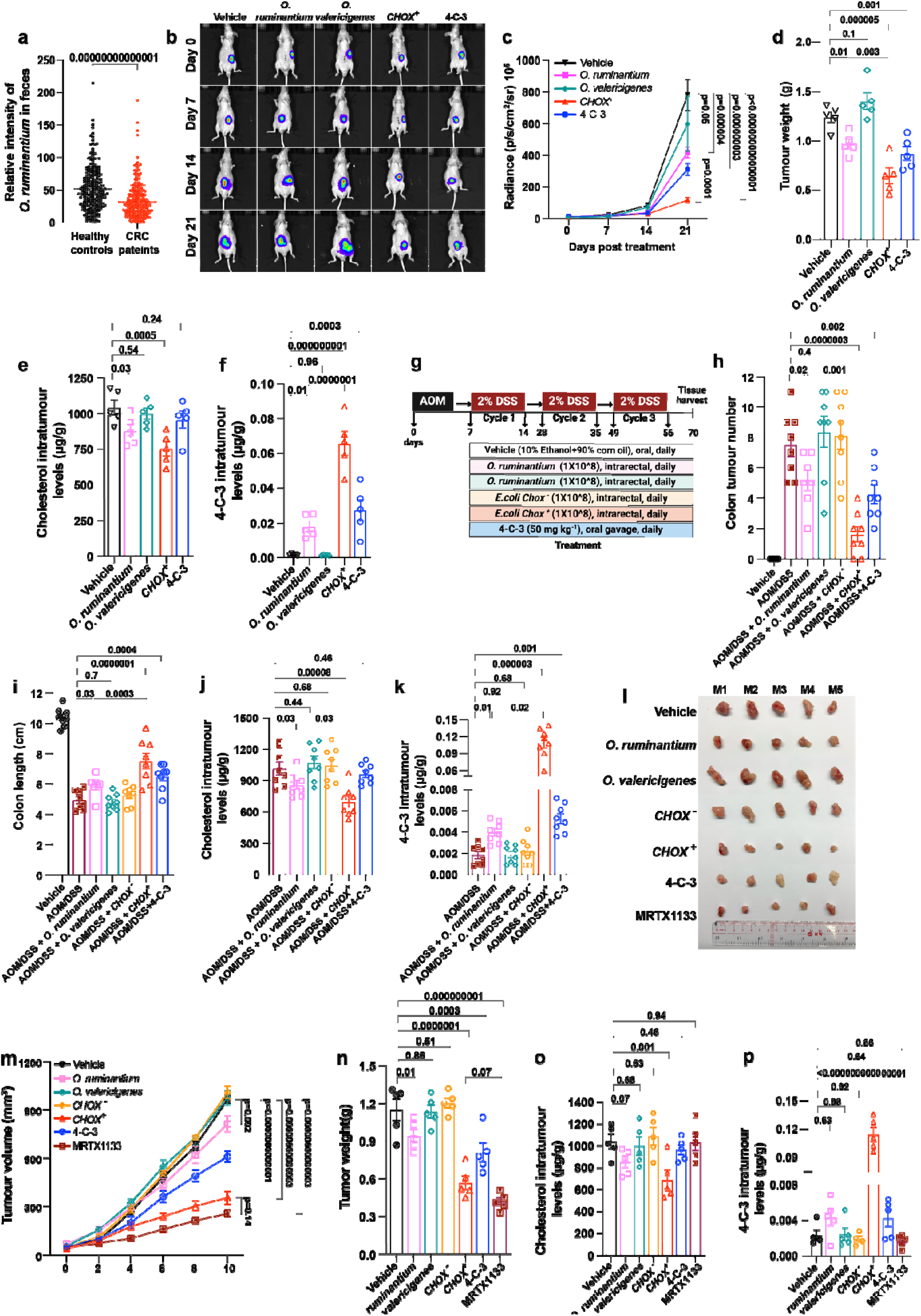
Suppression of colon tumorigensis by engineered microbe-mediated conversion of cholesterol to 4-cholesten-3-one (4-C-3) **(a)** The levels of *Ocillibacter ruminantium (O. ruminantium)* in patients with CRC and healthy individuals [mean ± s.e.m., n=532; unpaired t-tests]. **[b-f]** Orthotropic CRC model with luciferase-labelled HCT116 cells were subcutaneously implanted into nude mice to generate xenograft tumors. The mice were then treated daily with *O. ruminantium, O. valericigenes, E. coli CHOX^+^* bacteria or 4-C-3 for 21 days. **(b)** Images for luciferase signal of HCT116 xenograft tumors after the treatment. **(c)** Quantification of luciferase signal emitted by HCT116 xenograft tumors for the measurement of tumor growth. **(d)** Tumor weight. **(e-f)** The level of intratumoral cholesterol **(e)** and the level of intratumoral 4-C-3 **(f)**. [mean ± s.e.m., n=5 for each treatment group; two-way ANOVA for **(c)** & one-way ANOVA for **(d-f)**]. **[g-k]** C57BL/6 mice were treated with antibiotics and AOM-DSS regimen, followed by daily gavage with indicated treatment groups **(g)**. The colon tumor number **(h)**, colon length **(i)**, the level of intratumoral cholesterol **(j)**, and the level of intratumoral 4-C-3 **(k)** in the colon tissue of indicated treatment groups. [mean ± s.e.m., n=8 for each treatment group; one-way ANOVA for **(h-k)**]. **[l-p]** Immunocompetent mice were subcutanously implanted with CRC organoids from *Apc^min/+^Kras^G12D/+^*mutant mice and were intratumorally injected with *O. ruminantium, O. valericigenes, E. coli CHOX ^−^* bacteria, *E. coli CHOX^+^*bacteria, 4-C-3 or the KRAS^G12D^ inhibitor MRTX1133 for 10 days. **(l)** Images of tumors for indicated treatment groups are shown. **(m-p)** Tumor volume **(m)** and tumor weight **(n)**, the level of intratumoral cholesterol **(o)**, and the level of intratumoral 4-C-3 **(p).** [mean[±[s.e.m., n=5 for each treatment group; two-way ANOVA for **(m)** & one- way ANOVA for **(n-p)**].

## Discussion

High cholesterol levels are associated with an increased risk of CRC. Cholesterol-lowering strategies have potential for CRC prevention and treatment. However, therapeutic strategies for modulating cholesterol levels in CRC are limited and are usually associated with various complications, revealing an unmet clinical need for new therapeutic approaches for modulating cholesterol levels in the treatment of CRC. In this study, we introduce a new strategy in which the gut microbiome is leveraged to convert protumorigenic cholesterol into an anticancer metabolite; overall, the results revealed that engineered microbial therapies for restoring intratumoral cholesterol homeostasis constitute a safe and feasible therapeutic approach for the treatment of CRC. This study underscores the therapeutic potential of microbiome-mediated cholesterol metabolism in the treatment of CRC, providing comprehensive data on the efficacy of the cholesterol metabolite 4-C-3 in modulating CRC growth, and highlights a novel use for engineered bacteria in converting cholesterol into 4- C-3 to improve therapeutic outcomes.

To our knowledge, this study, for the first time, provides a proof-of-concept strategy, in which the microbiome converts intestinal cholesterol to its anticancer metabolites, that is a safe and feasible therapeutic approach for CRC. We also identified 4-C-3 as a novel antitumor microbial metabolite for CRC and delineated the previously unclear mechanism by which this metabolite exerts its anti-CRC effects. Furthermore, 4-C-3 does not affect noncancer cell lines, and mammalian cells do not have the capacity to convert cholesterol into 4-C-3, which makes our intervention with microbial engineering a unique and potentially valuable tool for treating CRC.

Our engineered microbes have several advantages over commercially available pharmacological agents in the control of cholesterol-mediated CRC tumorigenesis. Although accumulating evidence shows that statins, which are widely prescribed cholesterol-lowering agents that target cholesterol biosynthesis, have played a positive role in CRC prevention and treatment in recent decades, the clinical guidelines for the use of statins to treat CRC remain controversial(Cardwell *et al*., 2014; Emilsson *et al*., 2018; Liang *et al*., 2021; Voorneveld *et al*., 2017). Clinical trials have shown mixed results and the evidence is inconclusive (Lim et al., 2015), suggesting that controlling cholesterol levels alone may not be sufficient to control CRC development in clinical setting. The long-term use of statins is also associated with diverse complications (Khatiwada and Hong, 2024). In contrast, our engineered bacterial strain not only reduces intratumoral cholesterol levels but also has robust cancer-specific cytotoxic effects on CRC *in vivo* and *in vitro*. 4-C-3 is well tolerated *in vivo* at high doses, which reduces safety concerns in drug development. Furthermore, the manufacture of these synthetic biotic medicines is easily scalable, which is cost-effective in drug development. These results suggest that our intervention with microbial engineering for cholesterol metabolism represents an innovative and promising therapeutic approach for future clinical studies, as an adjunct to traditional therapy or as a preventive approach.

In this study, we identified 4-C-3 as a novel endogenous inhibitor of EGFR and revealed that its anti-CRC effect is primarily mediated via antagonism of the EGFR signaling pathway. As EGFR is one of the most common molecular targets in cancer therapy, particularly in CRC and other types of cancer, such as non-small cell lung cancer, the delivery of 4-C-3 via our unveiled microbiome strategy may have potential applications in other types of EGFR-related cancers. Indeed, our preliminary study revealed that 4-C-3 inhibits the growth of a panel of lung cancer cell lines (**Fig. S13)**. The therapeutic value of 4-C-3 is further supported by its ability to overcome resistance to common anti-EGFR therapies in CRC. Further investigations are needed to establish 4-C-3 as a promising therapeutic candidate at the pancancer level.

Our investigations revealed that 4-C-3 directly and negatively regulates EGFR via direct binding; thus, 4-C- 3 confers protection against CRC tumorigenesis via the suppression of EGFR signaling. These findings reveal a previously unrecognized mechanism for the control of EGFR signaling involving the microbial metabolism of cholesterol. Our new insights into the regulation of EGFR signaling may also provide a possible mechanism by which sensitivity to endogenous EGFR ligands is regulated in physiological and pathological situations, which deserves further investigation in the future. This study demonstrated that compared with potent EGFR ligands and EGFR-targeted therapeutics such as cetuximab, 4-C-3 binds to EGFR with relatively low affinity, suggesting that 4-C-3 may bind EGFR in a transient manner. Our computational simulation revealed that this transient interaction with the receptor may alter the conformation of EGFR and block the binding of EGFR ligands, thereby suppressing EGFR activation (**Fig. S6)**. Further investigations into the comprehensive mechanism of 4-C-3 in the regulation of EGFR signal transduction are needed.

EGFR-targeted therapies are crucial in treating colorectal cancer, especially in patients with metastatic CRC. These therapies work best in tumors without KRAS mutations, which can make tumors resistant to EGFR inhibitors. KRAS-targeted treatments are less effective against KRAS-mutant CRC partly due to EGFR-mediated reactivation of MAPK signaling. This challenge has led to clinical trials combining KRAS and EGFR inhibitors. However, this strategy faces limitations, as KRAS has been considered “undruggable” due to its strong affinity for GTP/GDP and lack of suitable binding pockets. Recent advances have produced KRAS^G12C^ inhibitors (e.g., sotorasib and adagrasib), but these do not target other KRAS mutations (e.g., KRAS p.G12D, G13D). In this study, we identified 4-C-3 as a novel compound that antagonizes both EGFR and oncogenic KRAS mutants. By targeting both EGFR and KRAS, 4-C-3 reduced resistance to anti-EGFR therapies caused by KRAS mutations (e.g., KRAS p.G12D) in CRC. This is significant because there are no FDA-approved drugs for KRAS^G12D^, which is more common in CRC than KRAS^G12C^. Additionally, 4-C-3 is effective against CRC cells with KRAS^G13D^, indicating its potential to target various KRAS mutants. Our findings show that 4-C-3 selectively inhibits KRAS^G12D^ more than wild-type KRAS, which is important for minimizing side effects and maximizing therapeutic efficacy. This selectivity is supported by the excellent safety profile of 4-C-3 in preclinical tests. Further research is needed to establish 4-C-3 as a promising pan-KRAS therapeutic. To our knowledge, this study is the first to identify a natural compound (4-C-3) that inhibits CRC tumorigenesis by targeting both EGFR and oncogenic KRAS mutants (e.g., KRAS^G12D^). Given its high selectivity for KRAS mutants over wild-type KRAS and its excellent safety profile, our study serves as a blueprint for developing safe and effective EGFR-KRAS therapies for CRC and other cancers driven by EGFR-KRAS. Further preclinical and clinical studies are necessary to validate these findings and assess the safety and efficacy of 4-C-3 in patients.

The idea that gut bacterial metabolism of cholesterol may control cholesterol levels was proposed over 100 years. However, the physiological function of this connection remains completely unknown. Despite the longstanding hypothesis, how these interactions might be leveraged for therapeutic purposes in conditions such as hypercholesterolemia and its associated cancers remains unexplored. Recent findings from observational studies in humans have shown that *Oscillibacter* can process cholesterol in the gut and is associated with lower cholesterol levels and reduced risks of pathological conditions, such as cardiovascular disease. Despite numerous correlation studies, experimental evidence showing the direct role of cholesterol-metabolizing bacteria in the regulation of cholesterol levels and cholesterol-related diseases is lacking. In this study, our proof-of-concept study revealed not only that the levels of 4-C-3 and its natural producer (*O. ruminantium*) are reduced in CRC patients but also that microbiome-mediated conversion of intestinal cholesterol to its anticancer metabolite 4-C-3 is a safe and feasible therapeutic approach for CRC. A detailed understanding of the microbes and metabolic pathways that affect CRC risk may eventually lead to personalized therapies that manipulate gut bacteria. This study integrates findings from human subjects with validation experiments in multiple animal models to provide actionable mechanistic insights that will serve as starting points to improve cancer care and treatment, potentially transforming the current therapeutic landscape.

There are limitations to this study. Although *O. ruminantium* has demonstrated the capacity of converting cholesterol into 4-C-3 and shown antitumor potency in multiple CRC models, these effects are weaker compared to those observed with *E. coli* CHOX+ bacteria. This suggests that while *O. ruminantium* has some potential in CRC treatment, its efficacy may be limited when used alone. It is therefore important to explore other bacterial species that are associated with altered cholesterol metabolism in CRC to identify those that might have stronger antitumor effects. Furthermore, investigating the potential synergistic effects of combining *O. ruminantium* with other bacterial species could provide insights into developing more effective bacterial-based therapies for CRC.

## Materials and Methods

### KEY RESOURCES TABLE

| REAGENT or RESOURCE | SOURCE | IDENTIFIER |
| --- | --- | --- |
| **Antibodies** |  |  |
| EREG (D4O5I) Rabbit mAb | Cell Signaling | Cat.#12048 |
| EGF Receptor (E746-A750del Specific) (D6B6) XP<sup>®</sup> Rabbit mAb | Cell Signaling | Cat. #2085S |
| Phospho-HER2/ErbB2 (Tyr1248)/EGFR (Tyr1173) Antibody | Cell Signaling | Cat. #2244 |
| Phospho-PI3 Kinase p85 (Tyr458)/p55 (Tyr199) (E3U1H) Rabbit mAb | Cell Signaling | Cat. #17366; |
| PI3 Kinase p110 $\delta$ (D1Q7R) Rabbit mAb | Cell Signaling | Cat. #34050 |
| mTOR (7C10) Rabbit mAb | Cell Signaling | Cat. #2983 |
| Phospho-mTOR (Ser2448) (D9C2) XP<sup>®</sup> Rabbit mAb | Cell Signaling | Cat.#5536 |
| Phospho-Akt (Ser473) (D9E) XP<sup>®</sup> Rabbit mAb | Cell Signaling | Cat.#4060 |
| K-Ras Antibody | Cell Signaling | Cat.#53270 |
| p44/42 MAPK (Erk1/2) | Cell Signaling | Cat.#4695 |
| Phospho-p44/42 MAPK (Erk1/2) | Cell Signaling | Cat.#9101 |
| Akt Antibody | Cell Signaling | Cat.#9272 |
| β-Actin (D6A8) Rabbit mAb | Cell Signaling | Cat.#8457 |
| **Bacterial strains** |  |  |
| *Oscillibacter ruminantium* | Deutsche Sammlung<br>von<br>Mikroorganismen<br>und Zellkulturen | DSM 113516 |
| Human feces | Clinical trials | N/A |
| **Chemicals and reagents** |  |  |
| Cholesterol | SIGMA | CSA.#57-88-5 |
| 5-Cholesten-3-one | SIGMA | CSA.#601-54-7 |
| 4-Cholesten-3-one | MCE | Cat.#HY-113365 |
| Coprostanone | SIGMA | CSA.#566-88-1 |
| Coprostanol | SIGMA | CSA.#360-68-9 |
| Dulbecco's modified Eagle's medium | Gibco | Cat# 11965 |
| Fetal bovine serum | Gibco | Cat# 26140 |
| LB Broth | Thermo Fisher | Cat# 12780052 |
| LB Agar | Thermo Fisher | Cat# 22700025 |
| RIPA buffer | Thermo Fisher | Cat #89900 |
| BCA Protein Assay Kit | Thermo Fisher | Cat #23225 |
| EASYpack Protease Inhibitor Cocktail | Roche | SKU# 5892970001 |
| PhosSTOP | Roche | SKU# 4906845001 |
| Enhanced chemiluminescence | Thermo Fisher | WBKLS0500 |
| TRIzol reagent | Invitrogen | N/A |
| One Step PrimeScript™ III RT-PCR Kit | TaKaRa | RR600A |
| Quantitect SYBR Green PCR Master Mix | Qiagen, Valencia, CA | N/A |
| **Cell lines** |  |  |
| HCT116 | ATCC | N/A |
| Colo205 | ATCC | N/A |
| HCT15 | ATCC | N/A |
| Cacao-2 | ATCC | N/A |
| DLD-1 | ATCC | N/A |
| HT55 | ATCC | N/A |
| SW620 | ATCC | N/A |
| YAMC | ATCC | N/A |
| LS513 | ATCC | N/A |
| MC38 | ATCC |  |
| **Plasmid** |  |  |
| EGFR G719S | Addgene | #11013 |
| pHAGE-EGFR-R222C | Addgene | #116293 |
| pHAGE-EGFR-S492R | Addgene | #116303 |
| pHAGE-EGFR-E114K | Addgene | #116238 |
| pHAGE-EGFR-P596L | Addgene | #116284 |
| **Recombinant human protein** |  |  |
| Human EGFR Protein, His Tag | DIMA Biotech | PME100099 |
| Amphiregulin (AREG) (NM_001657) Human Recombinant Protein | OriGene Technologies | TP761407 |
| Recombinant Human EREG protein (His tag) | Biosynth | 80R-4058 |
| Human KRAS / K-Ras (G12D & Q61H) Protein (His Tag) | Sino Biological | 12259-H07E1 |
| Human KRAS Protein, His Tag | DIMA Biotech | PME100649 |

#### Cell cultures

HCT116, Caco-2, SW620, HT55, HT29, Colo205, YAMC, DLD-1, LS513, MC38, and HCT15 cell lines were purchased from ATCC (Manassas, VA, USA). HCT116 cells with stable overexpression of firefly luciferase (Fluc) and enhanced green fluorescent protein (eGFP) were purchased from Imanis Life Sciences. All the cell lines were cultured in DMEM supplemented with 10% fetal bovine serum (FBS) and 1% penicillin/streptomycin a5% CO2 in a cell incubator.

#### MTT assay

Cell viability was determined by the 3-(4,5-dimethylthiazol-2-yl)-2,5-diphenyltetrazolium bromide (MTT) uptake method. Firstly, the cells (2,000-5,000 per well) were seeded in 96-well plates overnight prior to treatment. The optical density (O.D.) values for the control group were set as 100% viability.

#### Mouse models

The mice were purchased from the Animal Center of the Chinese University of Hong Kong and fed in the Laboratory Animal House of Hong Kong Baptist University. The mice were housed under a 12-hour light/dark cycle at a controlled temperature of approximately 25 and humidity (∼60%) with free access to food and water. The mice were fed standard laboratory chow diets for one week before starting the experiment to allow for adaptation. The procedures for all *in vivo* studies were approved by by the Committee on the Use of Human & Animal Subjects in Teaching & Research (HASC) at Hong Kong Baptist University and procedures were approved by the Department of Health under Hong Kong legislation.

Subcutaneous mouse models: six-week-old male BALB/c nude mice or C57BL/6 mice were subcutaneously inoculated with 5 × 10^6^ DLD-1 cells, 1 × 10^6^ HCT116 cells, 2 × 10^6^ LS516 cells, or 1 × 10^6^ MC38 cells in the left armpit. Once tumors were palpable (∼60 mm³), the tumor-bearing nude mice were randomly assigned for different treatments. At the end of the experiment, tumor dimensions were measured with a caliper, and tumor volume was calculated with the formula: (L×W^2^)/2, where L and W represent the length and width, respectively, with diameter in mm.

Orthotopic mouse models: athymic Balb/c mice aged 5–6 weeks were maintained in laminar-flow cabinets with free access to standard chow diets and sterile water. The orthotopic CRC tumor model was constructed as described. Briefly, mice were anesthetized and underwent laparotomy with the cecum exposed and exteriorized. HCT116-Fluc-Neo/GFP-puro cells (5 × 10 ) were suspended in 40 µl of PBS/Matrigel (1:1) and inoculated into the submucosa of the mice cecum with a 29-gauge needle. One week after cell inoculation, the mice were randomized into four groups. Mice in the control group were treated with vehicle, and mice in the treatment group were given 10 mg/kg, 25 mg/kg, and 50 mg/kg 4-C-3. Bioluminescence signal was monitored using the IVIS 2000 Imaging System coupled with Living Imaging Software (Caliper). D-luciferin substrate (150 mg/kg) was intraperitoneally injected into mice 10 minutes before the bioluminescence signal measurement. No blinding was done. Mice were sacrificed 24 days after the treatment, and the tumors were dissected and fixed in 4% paraformaldehyde for further analyses.

Patient-derived xenograft (PDX) models: Human colorectal tumor specimens (P=0) were collected from surgical patients at the First People’s Hospital of Huizhou. Samples were preserved during transit to the laboratory by immersion in 50ml centrifuge tubes containing saline with 10% penicillin-streptomycin and placed on ice. The cancer tissue specimens were thoroughly rinsed three times with a large amount of physiological saline containing 10% gentamicin sulfate to remove residual impurities from the tissue block. Tumor tissue was cut into pieces approximately 2 × 2 × 2 mm³ in size using a sterile scalpel and forceps. A puncture inoculation needle was utilized for subcutaneous inoculation, and the tumor tissue was implanted into the axillary area of the forelimb of a nude mouse. Then, the PDX model with growing P1 generation tumor cells was established. Tumor volume was calculated twice weekly using the formula: (L×W^2^)/2. When the P1 generation tumor grew to about 1000 mm³, the mice were euthanized, and the tumor surface mucous and connective tissues were removed under ice bath conditions. Next, the tumor mass was cut open, and the necrotic part was removed. The processed P1 generation tumor tissue was then inoculated into the axilla of 10 male nude mice aged 7–8 weeks and weighing 19 ± 1 g, according to the above tumor inoculation method. The formal experiment was conducted using PDX tumor tissues at P3 generation.

AOM/DSS models: Mice were randomly divided into four experimental groups and one control group (n = 8 per group). The experimental groups were intraperitoneally injected with a single dose of 10mg/kg azoxymethane (AOM) on the first day of the first week, then fed with drinking water supplemented with 2% Dextran Sodium Sulfate (DSS) for 1 week, followed by normal drinking water for 2 weeks, with 3 weeks as a cycle, which was repeated three times. In addition, mice in the 4-C-3 treatment group were orally administrated with 4-C-3 at the dosage of 50 mg/kg/day/mouse, and the treatment group with engineered bacteria were orally administrated with E. coli CHOX^+^/E. coli CHOX^-^ bacteria (1 × 10 CFU/day/mouse). In the control group, mice were treated with the same amount of solvent. At the end of the experiment, the mice were euthanized and the colons were opened longitudinally. The visible tumors were counted, and tumor dimensions were measured with a caliper. Tumor volume was calculated with the formula: (L×W^2^)/2. The tumor burden was calculated by summing the volumes of all tumors.

#### Human specimens

Fresh human colon cancer specimens were collected from CRC patients after surgical tumor resection at the First People’s Hospital of Huizhou. None of the patients had received chemotherapy or immunosuppressive therapy for at least 3 months prior to the surgery. To obtain the normal tissue, a distance of more than 5 cm to the tumors was kept. The use of tumor tissue samples for research purposes was approved by the Institutional Human Research Ethics Committee at the First People’s Hospital of Huizhou. This study was performed in accordance with the Declaration of Helsinki.

#### Patient-derived organoids

The isolation of patient-derived organoids was performed as our previously published study. Briefly, fresh human colon cancer specimens were collected from CRC patients after surgical tumor resection at the First People’s Hospital of Huizhou. The colonic tissues were washed thoroughly with ice-cold saline containing antibiotics and antifungals, and the underlying muscle layer was removed from the submucosal layer. The tissues were cut into fine pieces that were then incubated in a solution of collagenase and/or Dispase at 37°C with gentle agitation until the tissue is dissociated into single cells or small clusters. After isolation, cells were suspended in the matrix composed of advanced DMEM/F12 medium (Gibco) and growth-factor-reduced Matrigel in a ratio of 1:1. The cell-matrix mixture was seeded in 48-well plates (1000 single cells per 25 μl of Matrigel per well). The Matrigel was polymerized for 10 min at 37 °C and 250μl of the culture medium consisting of advanced DMEM/F12 medium + Glutamax (Invitrogen) containing N2, B27 supplements (Invitrogen), 10 mmol/L HEPES (Invitrogen), 1.25 mmol/L *N*-acetyl cysteine (Sigma-Aldrich), 2 mmol/L glutamine, 10μmol/L R-spondin-1, Noggin, WNT3A, SB202190 (Sigma-Aldrich), 50ng/mL EGF (Invitrogen), and 50μg/mL gentamycin. We developed five concentration gradients for the *in vitro* test. The PDO samples were treated with 4-C-3 for 5 days, and during the treatment period, the growth of the organoids was observed in a light microscope. At least 5 random fields were taken for quantification. The effect of the drugs was verified by observing the growth, activity, morphology, and cell clusters of the organoids.

#### Western blot analysis

The experiment was performed as described in our previous studies(Chow et al., 2022; Guo et al., 2024). After the treatments, the whole-cell lysates of the CRC cells were collected by suspending the cells in RIPA lysis buffer, then centrifuging the samples at 14,000 rpm for 10 minutes at 4 °C. The protein concentration was measured and calculated using a BCA Protein Assay Kit. Depending on the detection marker, 10 to 25μg of proteins were loaded into 10% SDS-PAGE and then transferred to PVDF membranes. After transfer, the membranes were incubated in blocking solution (5% skim milk powder in TBS containing Tween 20) for 1 hour at room temperature, and then the membranes were incubated with primary antibody overnight at 4 °C. After that, the membranes were incubated with secondary antibody for 1 hour at room temperature. All antibody solutions were diluted in TBS containing Tween 20 and 5% dry milk. The membranes were exposed using enhanced X-ray film and ECL reagent.

#### Real time PCR

The experiment was performed as described in our previous studies(Guo et al., 2022a; Guo et al., 2022b). Total RNA was extracted from samples using TRIzol reagent (Invitrogen) in accordance with the RNA extraction protocol of Thermo Fisher Scientific. cDNAs were subsequently prepared using the One Step PrimeScript™ III RT-PCR Kit (RR600A, TaKaRa). Real-time polymerase chain reaction (PCR) was performed using the Quantitect SYBR Green PCR Master Mix (Qiagen, Valencia, CA) with 1μL cDNA in a final volume of 10μL and the following primers at a final concentration of 1000nM. Primers for EREG were 5′-ACGTGTGGCTCAAGTGTCAA-3′ (forward) and 5′-CACTTCACACCTGCAGTAGTTT-3′ (reverse).

Amplification was performed using the LightCycler 2000 instrument (Roche, Indianapolis, IN). The cycling conditions comprised a denaturation step for 15 minutes at 95 °C, followed by 40 cycles of denaturation (95 °C for 15 seconds), annealing (59 °C for 20 seconds), and extension (72 °C for 15 seconds). After amplification, a melting curve analysis was performed with denaturation at 95 °C for 5 seconds, then continuous fluorescence measurement was made from 70 °C to 95 °C at 0.1 °C/second.

#### Flow cytometer analysis

Flow cytometry analysis was performed to determine cell apoptosis. Treated cells were harvested using trypsin without EDTA and phenol red to prevent any interference with the staining process. After harvesting, the cells were washed twice with cold PBS to remove any residual trypsin and other contaminants. Subsequently, 100μL of the cell suspension was transferred to a 5 mL culture tube. Cells were then incubated with 5μL of FITC-conjugated annexin-V reagent (2.5 mg/mL) and 5 μL of propidium iodide (PI) solution (5 mg/mL) for 15 minutes at room temperature in the dark. After incubation, 400μL of binding buffer was added to each tube. The stained cells were immediately analyzed using a flow cytometer equipped with appropriate filters for FITC and PI detection. Data was acquired and analyzed using FlowJo software to determine the percentage of apoptotic cells.

#### Human study

522 fecal samples were collected from the Fudan University Shanghai Cancer Center, Shanghai, China, and the Second Hospital of Shandong University, Shandong, China, from 2018 to 2021. Patients were diagnosed with CRC following postoperative pathological examination of tissue biopsies collected during colonoscopy. Fecal samples were obtained from patients at the hospital approximately two weeks after colonoscopy and stored at -80°C before undergoing metagenomic sequencing and metabolomics profiling. Participants’ demographics and clinicopathological characteristics, including age, gender, tumor location, size, differentiation, TNM stage, KRAS/NRAS/BRAF mutation, nerve invasion, lymphatic invasion, vascular invasion, MMR status, administration of neoadjuvant therapy, administration of adjuvant therapy, history of diabetes, hypertension, and other cardiovascular diseases, were collected from the electronic medical record system. The healthy controls were recruited volunteers with no gastrointestinal tumors confirmed by colonoscopy screening. Healthy controls were excluded if they had a history of enteric disorders including chronic diarrhea, inflammatory bowel disease, or gastroesophageal reflux disease; previous cardiovascular disease; diabetes mellitus; previous other malignancies; antibiotic use within six months before enrollment; or other infectious diseases.

#### Mass spectrometry analysis

An AGILENT ultra-performance liquid chromatography (UPLC) system coupled to a triple quadrupole (QQQ) MS6460 mass spectrometer was used for the targeted metabolomics profiling study. A Waters BEH 2.1x50 mm C18 1.7μm column with a pre-column was used. The mobile phase used in LC-MS-QQQ was A: water and B: MeOH. The gradients were set as 80% B (0-10 min), 100% B (10-12 min), 80% B (12-12.1 min), 80% B (12.1-14 min). In targeted metabolomics, the MS data was collected and processed by in-house software provided by Agilent.

Approximately 80 mg of tumor tissue or adjacent normal tissue samples were mixed with 1 mL of methanol after adding steel beads. The mixture was vigorously vortexed and then incubated overnight at 4°C. Following this, the mixture was centrifuged at 15,000 rpm for 10 minutes at 4°C. The supernatant (200 µL) was collected for LC-MS analysis.

#### Bacteria strains

*Oscillibacter ruminantium (*DSM113516) were cultured in Medium 104b under anaerobic conditions at 37°C. The Chox gene was cloned into the pUC57 vector using standard molecular cloning techniques, including restriction digestion and ligation. The resulting plasmid was transformed into E. coli (TOP10) by heat shock transformation. The insertion of the Chox gene into E. coli was confirmed by PCR using gene-specific primers and agarose gel electrophoresis. Additionally, a pUC57 vector-only control strain of E. coli was constructed by transforming E. coli with the empty pUC57 vector. Both the **Chox**-expressing **E. coli** and the vector-only control strains were incubated in LB medium supplemented with 100 μg/mL ampicillin and 0.5% cholesterol at 37°C for 24 hours with shaking at 200 rpm to ensure proper aeration and nutrient distribution. Post-incubation, the culture broth was centrifuged at 5,000 r.p.m. for 10 minutes to pellet the bacterial cells. The supernatant was collected, and the cholesterol and its metabolites in the culture broth were extracted using a liquid-liquid extraction method with chloroform and methanol. The extracted metabolites were then dried under nitrogen gas, reconstituted in a suitable solvent (e.g., methanol), and quantified using LC-MS. The LC-MS analysis was performed on an Agilent 1290 Infinity II coupled with an Agilent 6460 Triple Quadrupole mass spectrometer, using a C18 reverse-phase column (2.1x50 mm, 1.7 μm particle size) with a binary solvent system consisting of water (solvent A) and acetonitrile (solvent B) containing 0.1% formic acid. The gradient elution was set from 10% B to 90% B over 15 minutes, with a flow rate of 0.3 mL/min. Quantification was achieved by comparing the retention times and mass spectra of the samples to those of cholesterol and its known metabolites.

#### Cellular thermal shift assay

Cells treated with 4-C-3 were disrupted by liquid nitrogen, then centrifuged at 12,000 r.p.m. for 15 min to obtain total protein extracts. The proteins were divided into ten fractions and denatured at different temperatures for 10 min. Denatured proteins were precipitated by centrifugation at 12,000 r.p.m. for 10 min. EGFR and KRAS were detected by Western blot.

#### Microscale thermophoresis

Recombinant human protein was labeled with a Monolith NT Protein Labeling Kit RED-NHS (Nanotemper, Munich, Germany) and diluted to 50 nM with HEPES buffer (10 mM HEPES, 150 mM NaCl, 0.05% Tween). 4C3 was diluted with 1% ethanol in HEPES buffer to concentrations ranging from 125 μM to 3.8 nM. Samples were incubated at room temperature for 5 minutes. The mixtures were centrifuged at 16,000 × g for 5 minutes and loaded into capillaries. Microscale thermophoresis measurements were performed using a Monolith NT.115 (Nanotemper, Munich, Germany).

#### Ras activation assay

The Ras activation assay was conducted using a Pan-Ras Activation Kit (Cell Biolabs) following the manufacturer’s protocol. For control samples, cell lysates were treated with GTPγS and GDP at 30°C for 30 minutes, then MgCl was added. The lysates were then mixed with Raf1-RBD GST-conjugated agarose beads for 1 hour at 4°C, followed by washing. The proteins bound to Raf1-RBD were eluted using 2× sample loading buffer, and the co-precipitated Ras protein was analyzed via Western blotting.

#### Molecular dynamics simulation

Molecular dynamics simulations were conducted using Schrödinger software and the OPLS 2005 force field. The initial dimensions of the periodic box were 10 nm x 10 nm x 10 nm. The necessary number of counter ions (sodium cations) were added to ensure the electric neutrality of the system. The simulation was performed with periodic boundary conditions in all three directions, using a 100 ps timestep. The energy of the system was initially minimized using the steepest descent algorithm. Equilibrations were carried out in two steps. First, an NVT (constant Number of atoms, Volume, and Temperature) simulation was performed to bring the system to the target temperature. Second, an NPT (constant Number of atoms, Pressure, and Temperature) simulation was performed. Each ensemble was used to stabilize the temperature and pressure at 300 K and 1.01 bar, respectively. During the production dynamics, the structure was simulated for 100ps using the NPT ensemble. Finally, the simulation was run for 30ns.

#### Docking

The molecular docking was performed using the Glide module in Schrödinger. The binding site was selected as the center of the internal ligand in the crystal structure. The docking was performed with standard precision (SP-docking), and the GlideScore built into Schrödinger was used as the scoring function. For each docking simulation, 10 conformations were generated for each small molecule, and their energies were minimized after docking. For each conformation generated by docking small molecules, the low-energy binding conformation was selected for analysis.

#### Quantification and statistical analysis

All the results were obtained from multiple experiments (at least three independent experiments). Data were expressed as average with SD or SEM values where appropriate. Significance p-values were calculated using GraphPad Prism 8, and p-values less than 0.05 were regarded as statistically significant. Spearman’s rank coefficient correlation analysis was used for metabolite quantity correlation and the correlation between cholesterol and bacteria derived from patients with CRC. The Wilcoxon rank-sum two-tailed test was used to determine metabolite differences between CRC and healthy participants. The Wilcoxon one-tailed test was used to determine whether CRC-associated gut microbiota produces lower levels of cholesterol metabolites in the metagenomic data. Unpaired t-tests or one-way ANOVA were used in other experiments as indicated in the figure legends.

## Acknowledgements

The presented work was kindly supported by General Research Fund (12102020; W.H.L.X.), Health and Medical Research Fund (08793626; W.H.L.X.), Innovation and Technology Commission (ITS/058/22MS; W.H.L.X.), National Natural Science Fund (32322091; W.H.L.X.), Guangdong Medical University Research Foundation (4SG24205G; H. X.); Natural Science Foundation of Guangdong Province of China (2023A1515011116; H. X.); Discipline construction project of Guangdong Medical University (4SG23282G; H. X.); Dongguan Science and Technology of Social Development Program (20231800939832; H. X.); Natural Science Foundation of Guangdong Province (2022A1515011302; H. X.); National Natural Science Foundation of China (82070535 & 82370601; R. Y.); Natural Science Foundation of Sichuan Province (2022NSFSC0706; R. Y.); Bureau of Science and Technology Nanchong City (20SXQT0312; R. Y.) and Natural Science Foundation of Sichuan Province (MZGC20230048; R. Y.).

