## Supplementary figure for "Metabolic modulation of intratumoral cholesterol with gut microbiota for the treatment of colorectal cancer"

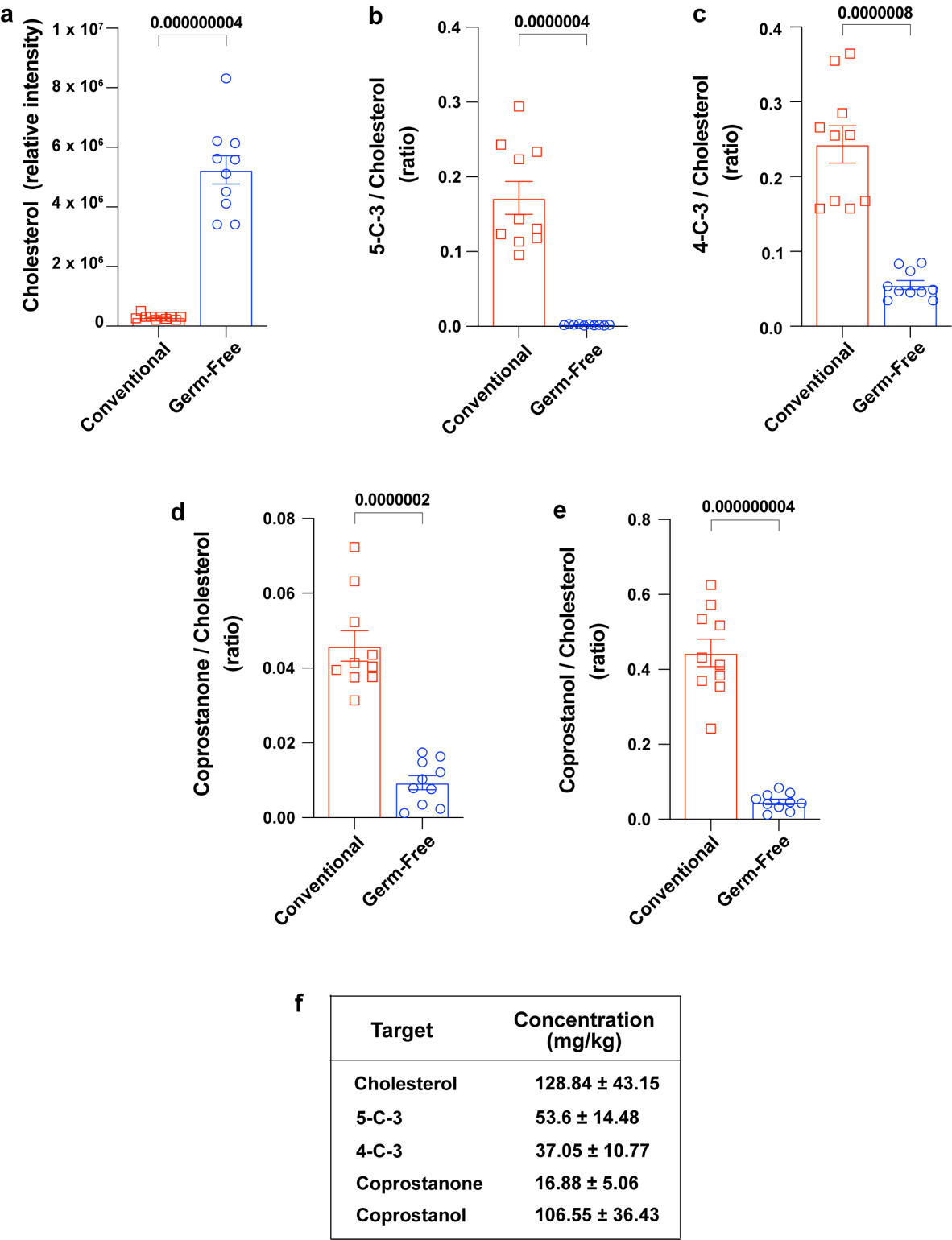

**Figure S1. The level of cholesterol and its microbial metabolites in germ-free mice and healthy individuals**

**(a-e)** The relative level of cholesterol and its indicated microbial metabolites in germ-free mice and conventional mice. [mean ± s.e.m., n=10 in each group; unpaired t-tests] **(f)** The concentration of cholesterol and its indicated microbial metabolites in healthy individuals [mean ± s.e.m., n=7]

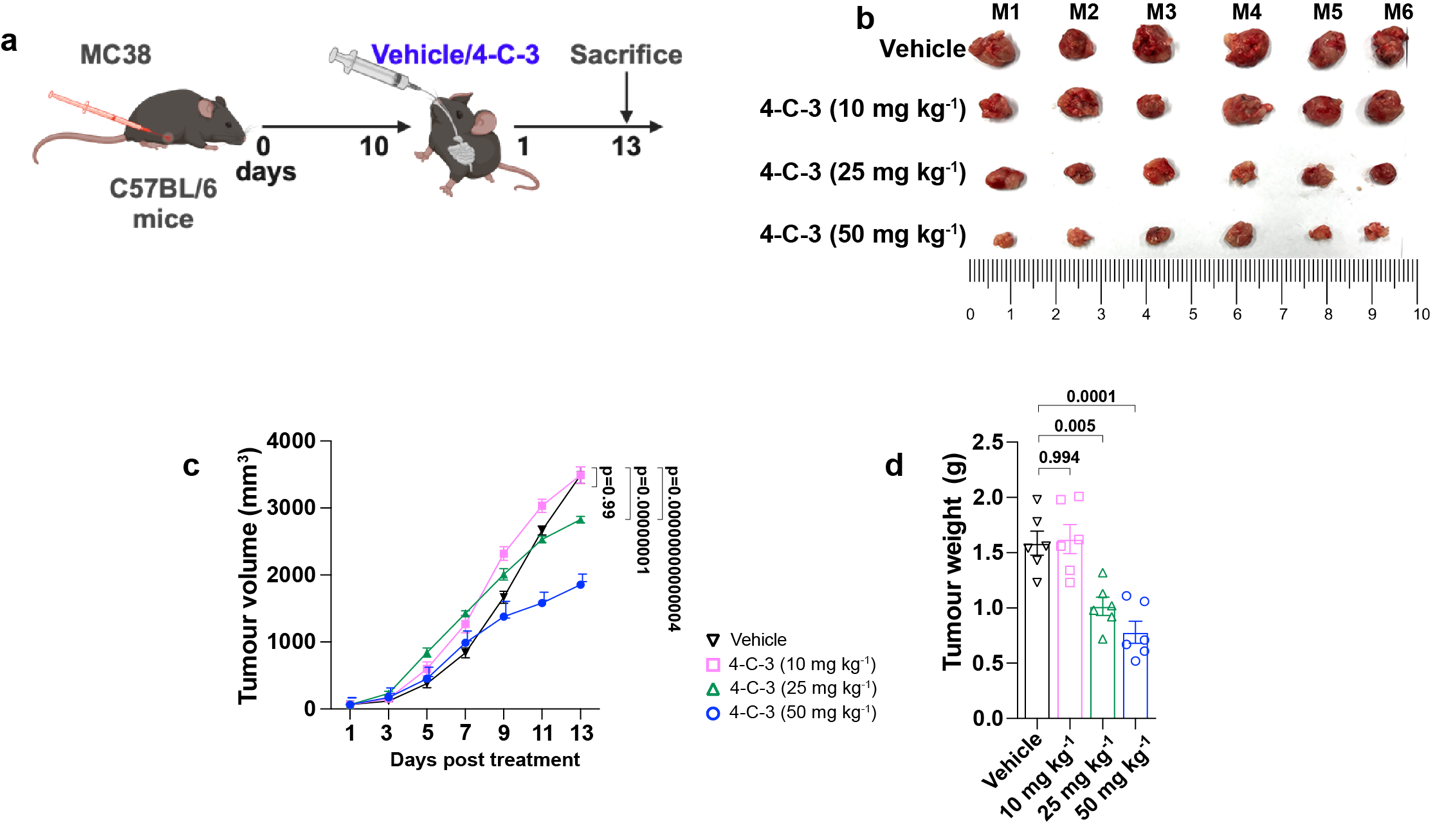

**Figure S2. 4-cholesten-3-one (4-C-3) suppresses colon tumorigenesis *in vivo***

**(a)** Schematic diagram showing the experimental design for establishing subcutaneous CRC model with MC38 cells and the treatment with 4-C-3 at indicated dosages. **(b)** Images of MC38 xenograft tumors after the treatment with 4-C-3. Tumor weight **(c)** and tumor volume **(d)** of MC38 xenograft tumors of mice in the indicated groups. [mean ± s.e.m., n=6 for each treatment group; two-way ANOVA for **(c)** & one-way ANOVA for **(d)**]. **(related to Fig. 2)**

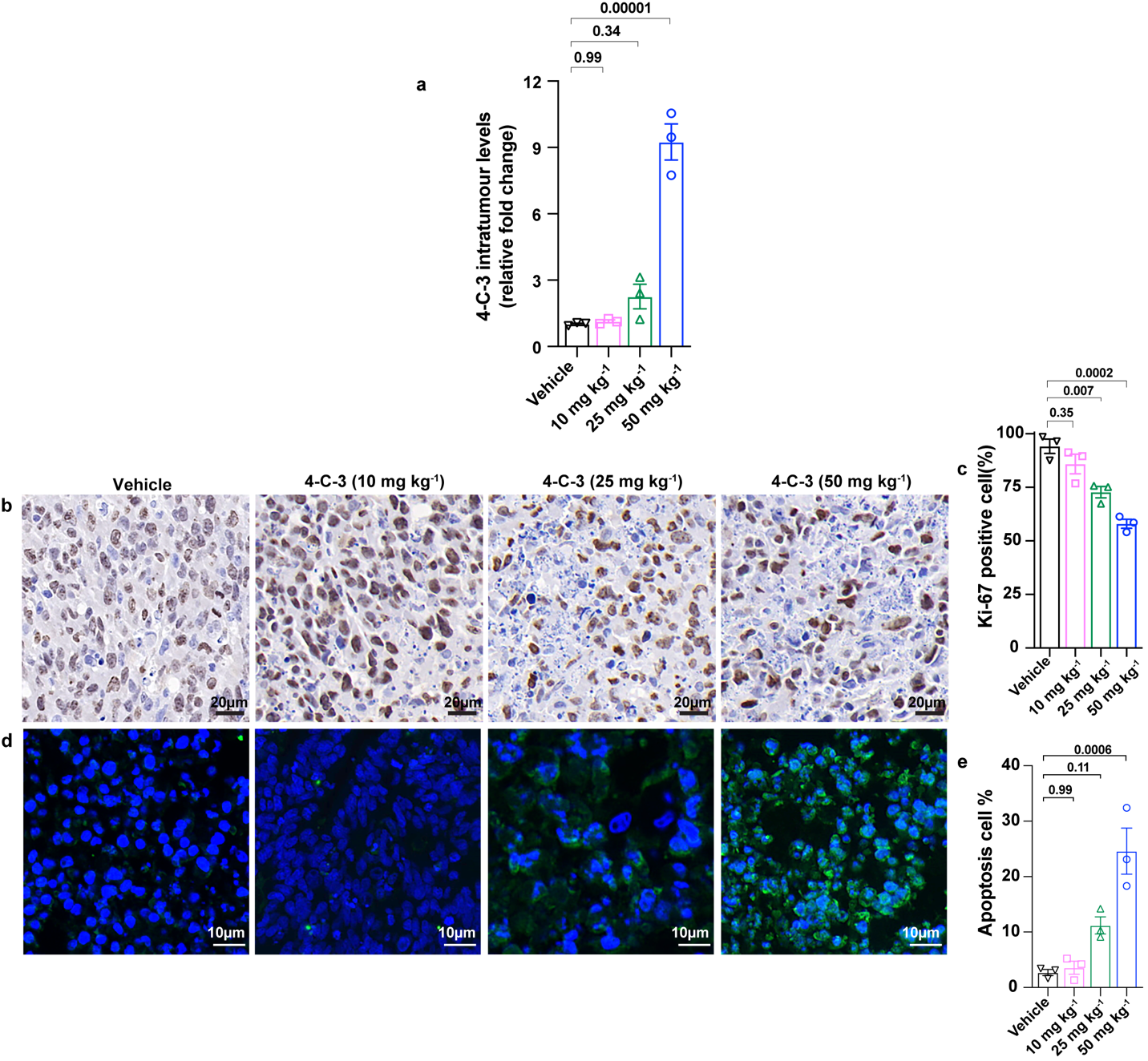

**Figure S3. The antitumor effect of 4-cholesten-3-one (4-C-3) on orthoptic xenografts of CRC cells**

**(a-e)** The level of intratumoral 4-C-3 **(a)**, the proliferation of cancer cells assessed by immunostaining of Ki-67 **(b-c)** and the apoptosis of cancer cells by TUNEL assay **(d-e)** in orthoptic xenografts of HCT116 cells upon 4-C-3 treatment. [mean ± s.e.m., n=3 for each treatment group; one-way ANOVA for a, c & e] **(related to Fig. 2)**

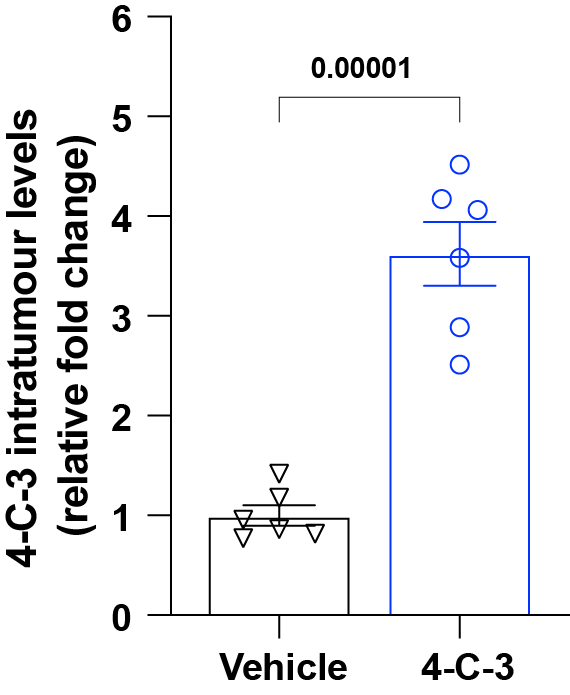

**Figure S4.** The intratumoral 4-C-3 levels in patient-derived xenografts (PDX) after treatment. [mean ± s.e.m., n=6 for each treatment group; unpaired t-tests].

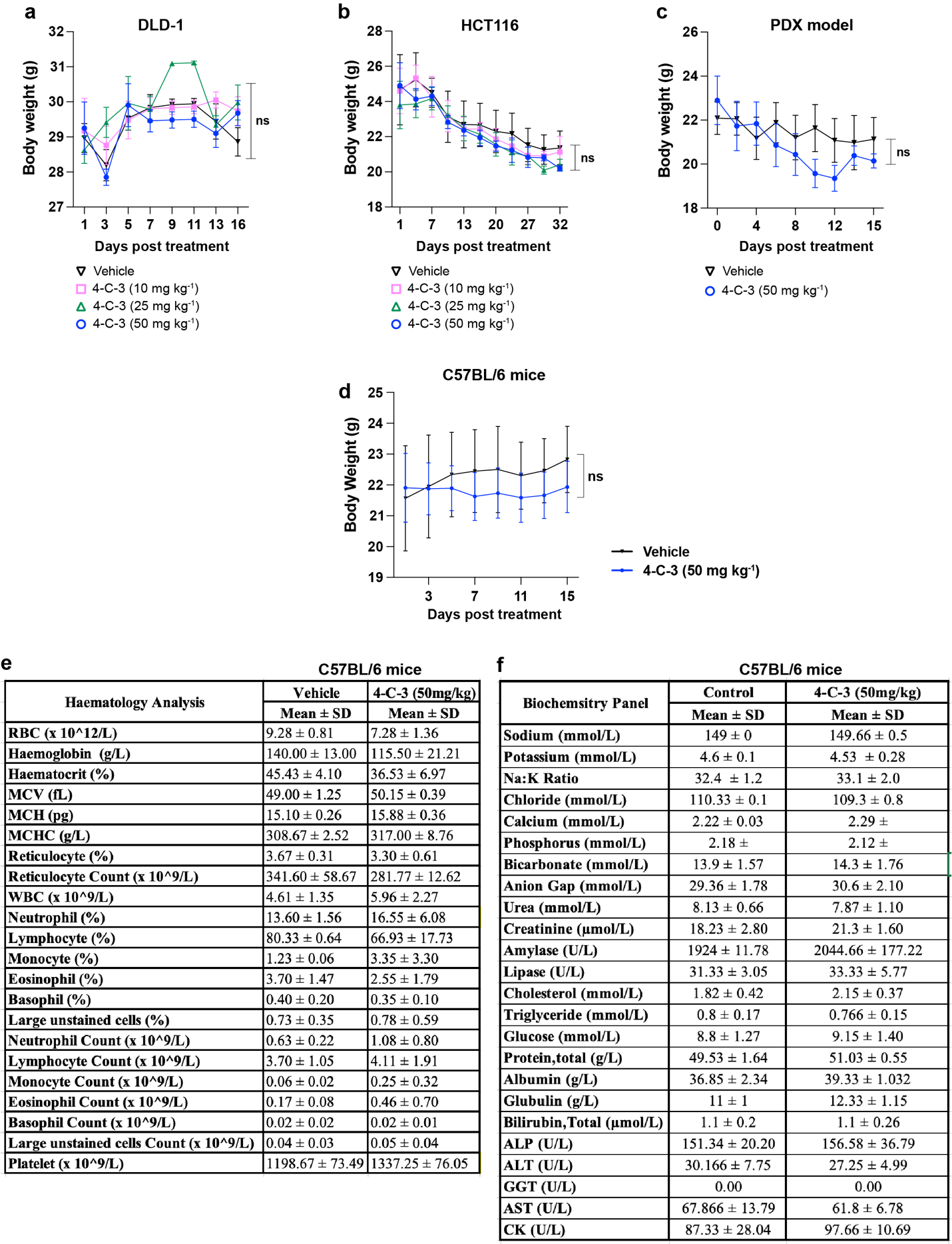

**Figure S5. The safety profile of 4-cholesten-3-one (4-C-3) in mice**

**(a-c)** The changes in body weight in mice bearing DLD-1 xenograft tumors **(a)**, orthoptic HCT116 xenografts **(b)**, and patient derived xenograft (PDX) tumors **(c)** with the treatment of 4-C-3 at indicated dosages [mean ± s.e.m., n=5 for (a), n=3 for (b), and n=6 for (c); two-way ANOVA]. **(d-f)** C57BL mice at the age of 8 weeks were treated with 4-C-3 (50mg/kg) for 4 weeks. The changes in body weight in mice were shown in **(d).**  The blood samples were collected for regular blood tests and different parameters were shown in **(e-f)** [mean ± SD, n=4 for each treatment group]

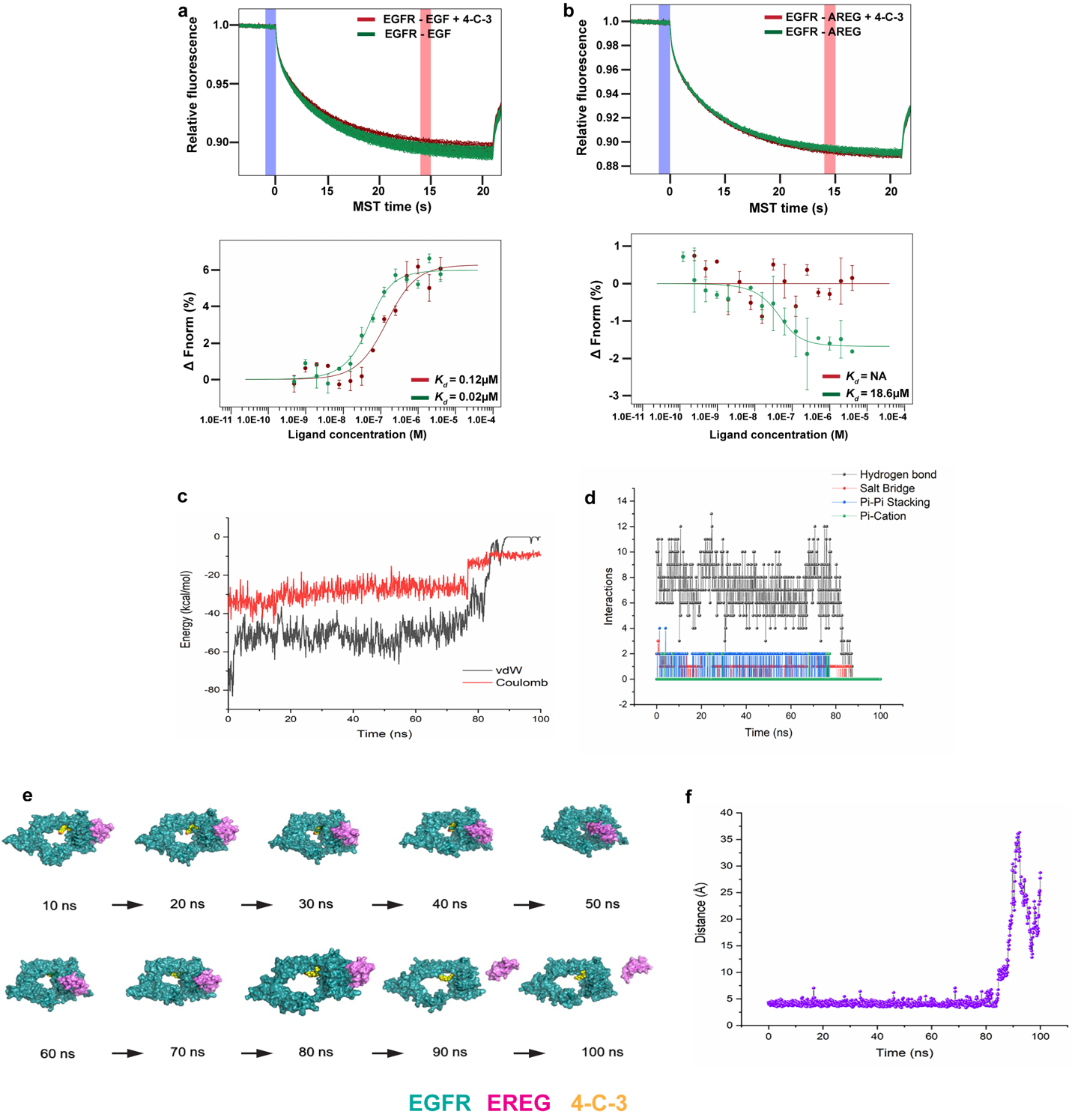

**Figure S6. The blockage of ligand binding of EGFR by 4-cholesten-3-one (4-C-3)**

**(a-b)** Microscale thermophoresis (MST) analyses for the binding between the recombinant extracellular domain of EGFR (EGFR-ECD) and recombinant EGF **(a)** and the binding between EGFR-ECD and recombinant AREG **(b)** in the presence/absence of 4-C-3. (n-3) **(c-f)** 100ns molecular dynamics (MD) simulations coupled with mechanics and thermodynamic calculations to study the dynamical structural characteristics and interactions between EGFR and EREG in the presence of 4-C-3.

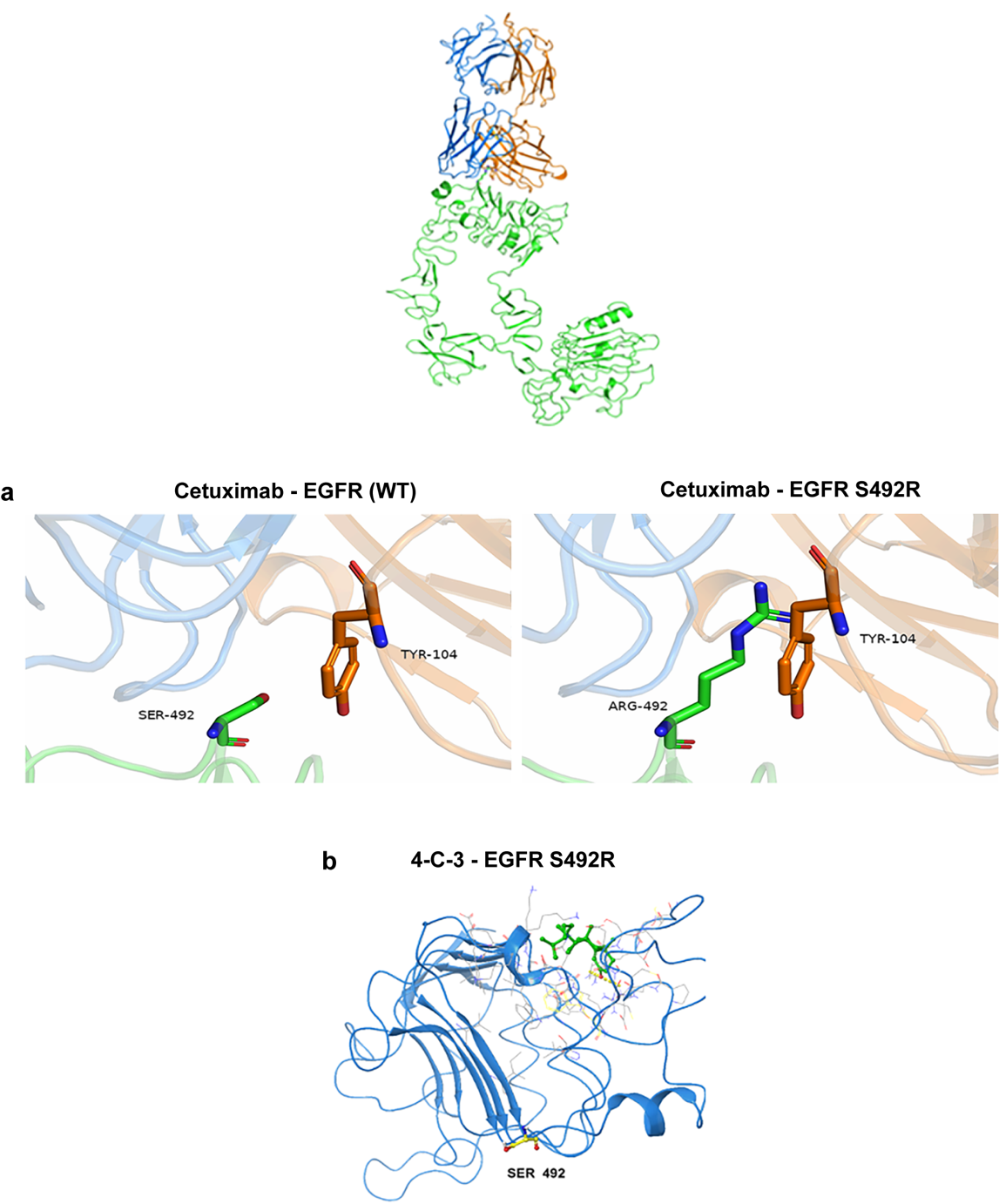

**Figure S7. The binding of cetuximab to EGFR variants**

**(a-b)** Molecular docking analyses of the binding between EGFR^WT^/EGFR^S492R^ and cetuximab **(a)** and the binding between EGFR^S492R^ and 4-C-3 **(b)**

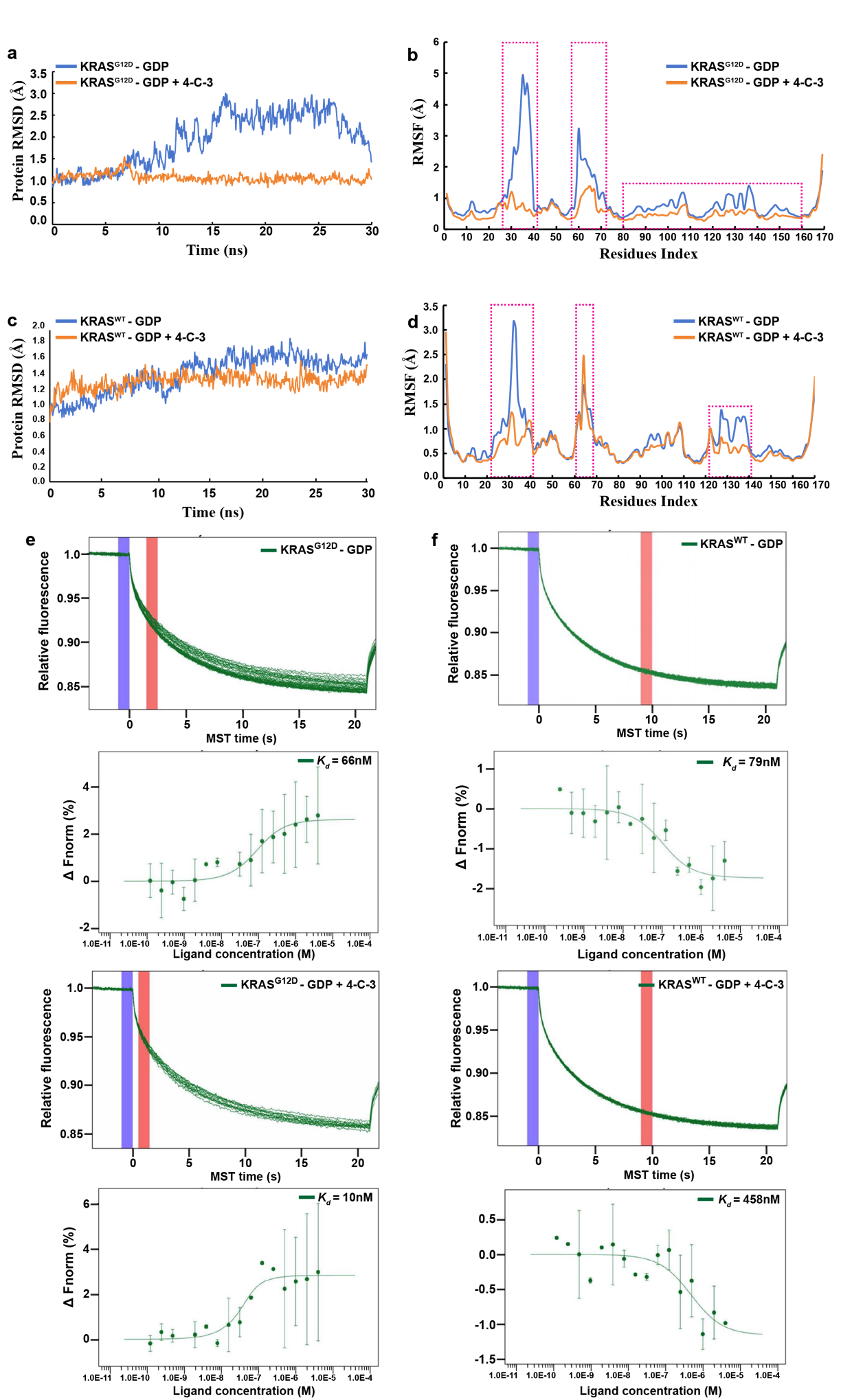

**Figure S8. Computational stimulation for the binding of 4-C-3 to oncogenic KRAS variants**

**(a-d)** Long timescale simulations were performed to reveal the interaction dynamics of the 4-C-3-KRAS^G12D^ complex**.** The analyses of the protein backbone root-mean-square fluctuation (RMSF) for the 4-C-3-KRAS^G12D^ complex **(a-b)** and the 4-C-3-KRAS^WT^ complex **(c-d). (e-f)** Microscale thermophoresis (MST) analyses for the binding of GDP on KRAS^G12D^ **(e)** and KRAS^WT^ in the presence/absence of 4-C-3 **(f)**.

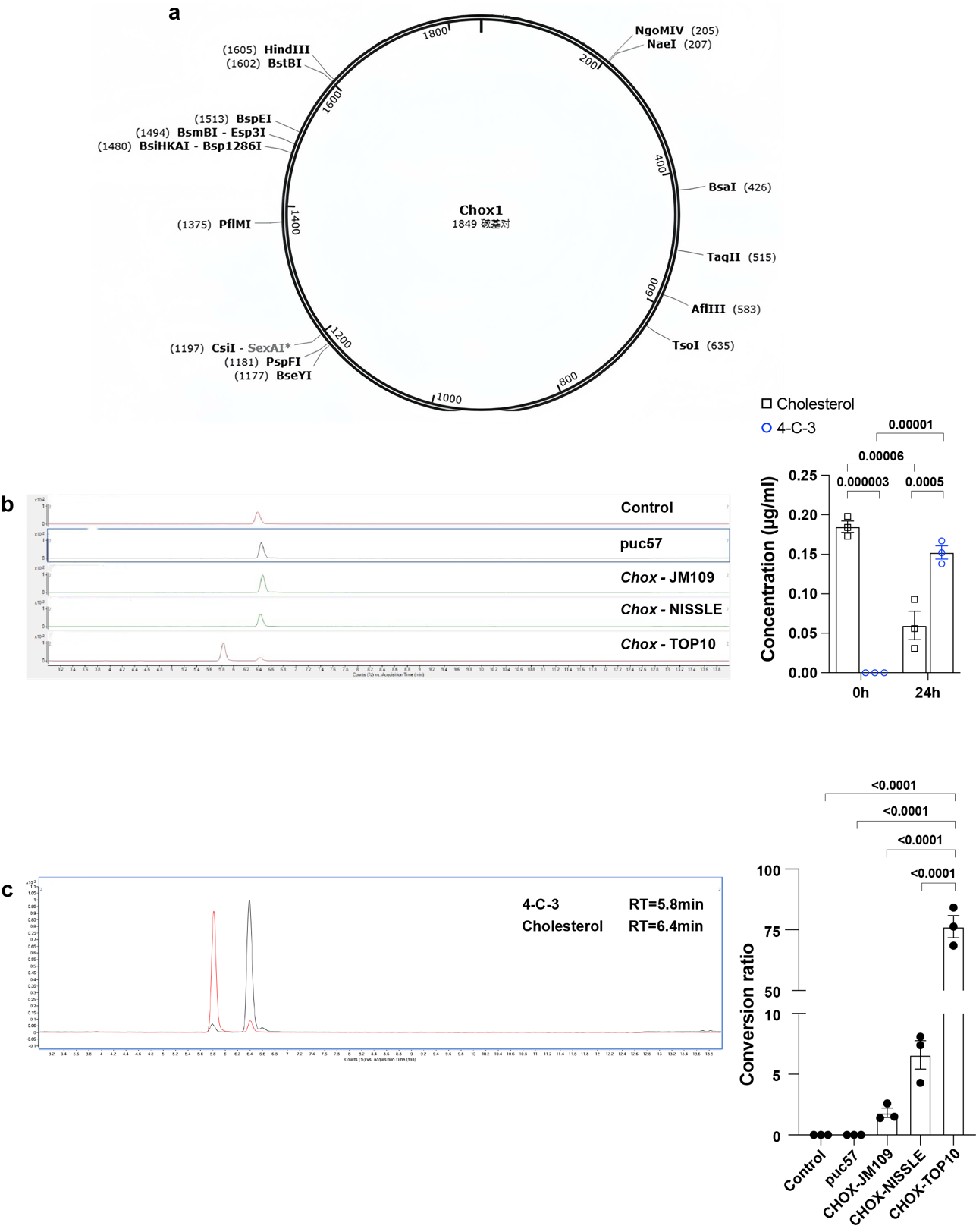

**Figure S9. Construction of engineered microbes for the conversion of cholesterol into 4-cholesten-3-one (4-C-3)**

**(a)** The construction map for the construct for expressing Chox1 **(b-c)** The *in vitro* conversion of cholesterol into 4-C-3 by indicated engineered bacteria including *E. coli*. (TOP 10), *E. coli*. (JM109) and *E. coli*. (Nissle 1917), as assessed by LC-MS analyses. [mean ± s.e.m., n=3 for each treatment group; one-way ANOVA]

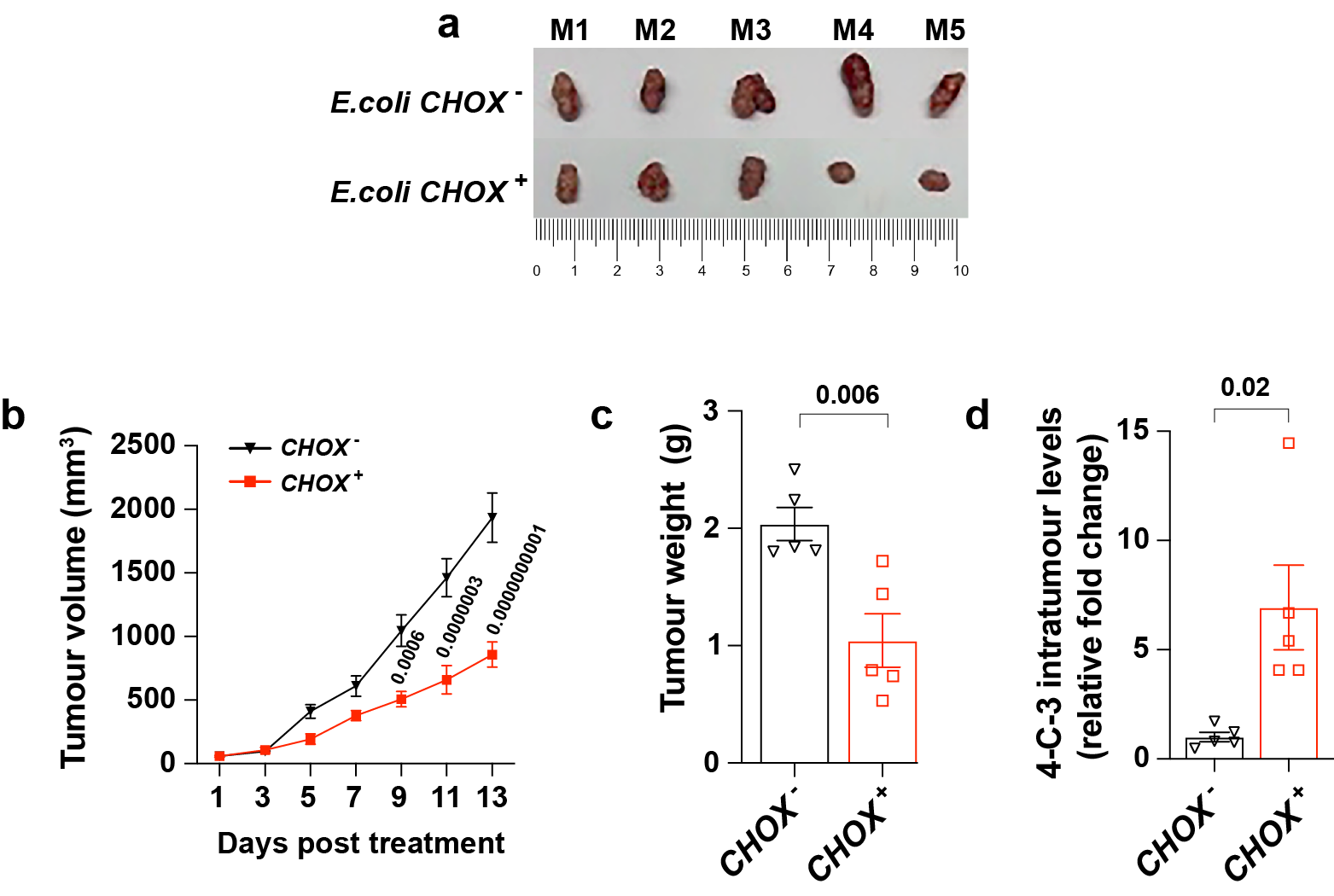

**Figure S10. Anti-tumour activity of engineered *E. coli CHOX^+^* bacteria in producing 4-cholesten-3-one (4-C-3)**

**[a-d]** MC38 colon cancer cells were subcutaneously implanted into nude mice to generate xenograft tumors. The mice were then gavaged daily with *E. coli CHOX^+^* bacteria or *E. coli CHOX^-^* bacteria for 13 days. Images of MC38 xenografts for indicated treatment groups are shown in **(a)**. **(b-d)** Tumor volume **(b)**, tumor weight **(c)** and the level of intratumoral 4-C-3 **(d)** of MC38 xenografts for indicated treatment groups. [mean ± s.e.m., n=5 for each treatment group; two-way ANOVA for **(b)**, unpaired t-test for **(c & d)**]

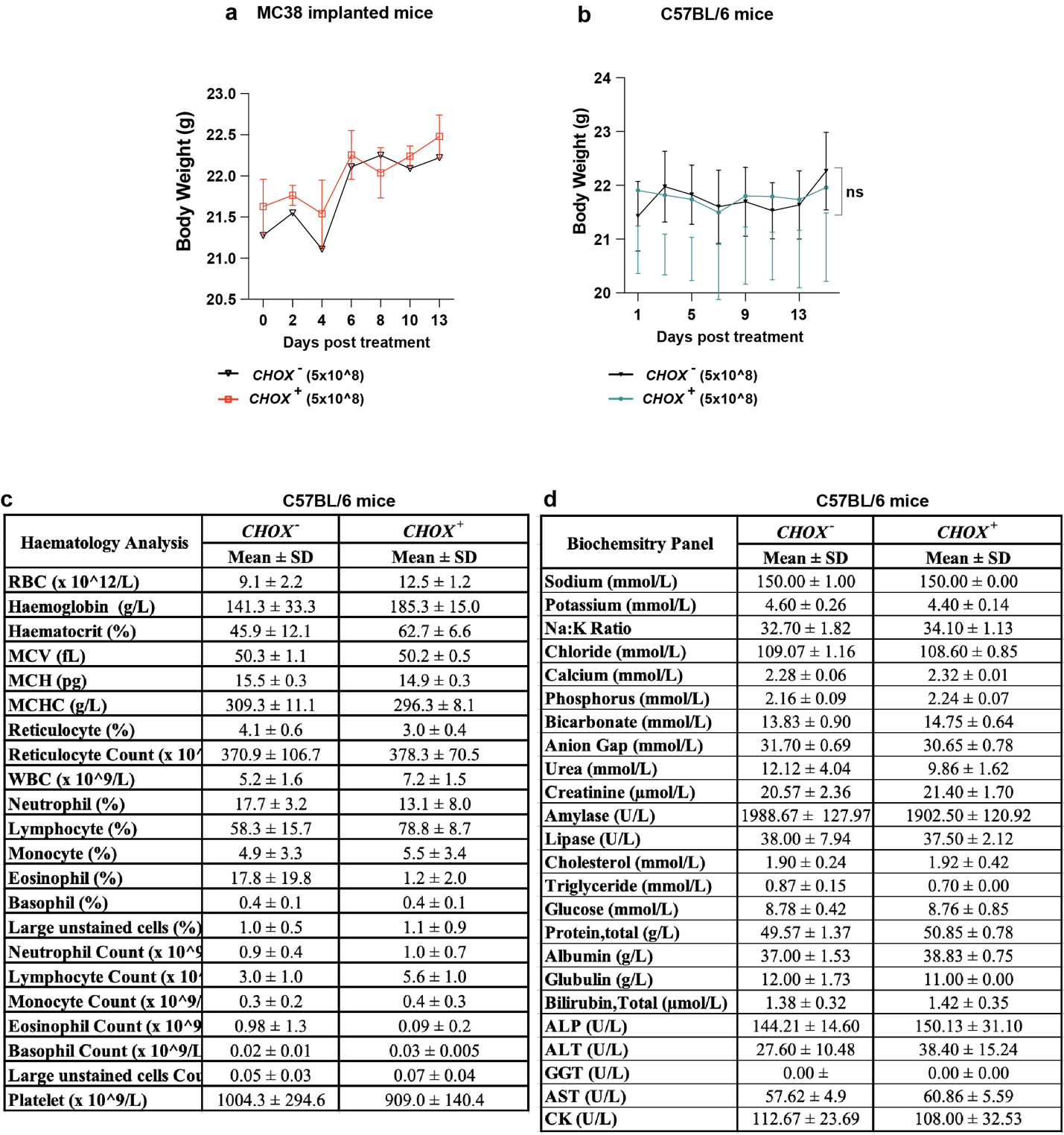

**Figure S11. The safety profile of *E. coli CHOX^+^* bacteria in mice**

**(a)** The changes in body weight of orthoptic HCT116 xenografts implanted mice upon the treatment *E. coli CHOX^+^* bacteria or *E. coli CHOX^-^* bacteria for 13 days [mean ± s.e.m.; n=5; two-way ANOVA]. **(b-d)** C57BL mice at the age of 8 weeks were daily gavaged with *E. coli CHOX^+^* or *E. coli CHOX^-^* bacteria (5X10^8^) for 4 weeks. The changes in body weight in mice were shown in **(b)**. The blood samples were collected for regular blood tests for haematological **(c)** and biochemical analyses **(d)** [mean ± SD, n=4 for each treatment group]

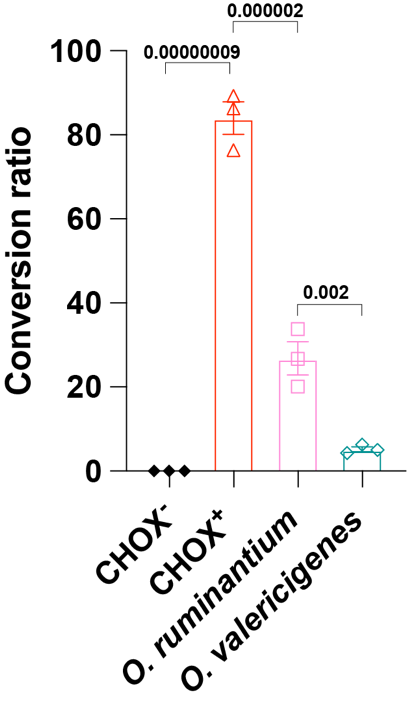

**Figure S12. The efficacy of *E. coli CHOX^+^* bacteria for converting cholesterol into 4-cholesten-3-one (4-C-3)**

The *in vitro* conversion of cholesterol into 4-C-3 by *E. coli CHOX^-^, E. coli CHOX^+^, Oscillibacter ruminantium and Oscillibacter valericigenes* bacteria as assessed by LC-MS analyses. [mean ± s.e.m., n=3 for each treatment group; one-way ANOVA]

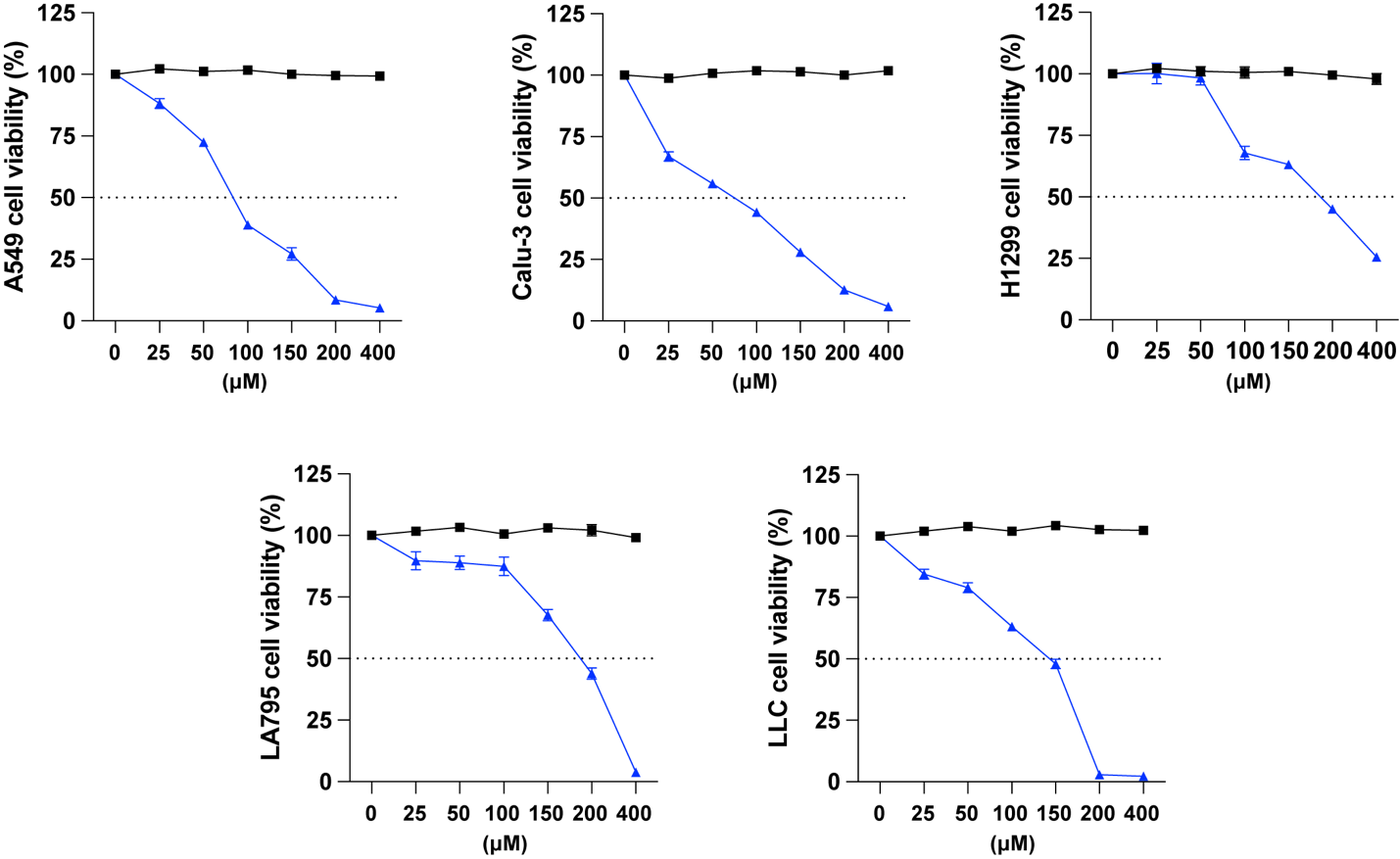

**Figure S13. Anti-tumor activity of 4-cholesten-3-one (4-C-3) in human lung cancer cells**

MTT assay in a panel of human colorectal cancer cell lines (A549, Calu-3, HT299, LA795, DLD-1 and LLC) treated with 4-C-3 at indicated dosages for 48 h. (mean ± s.e.m., n=3)
